# Comparison of spatial transcriptomics technologies using tumor cryosections

**DOI:** 10.1101/2024.04.03.586404

**Authors:** Anne Rademacher, Alik Huseynov, Michele Bortolomeazzi, Sina Jasmin Wille, Sabrina Schumacher, Pooja Sant, Denise Keitel, Konstantin Okonechnikov, David R. Ghasemi, Kristian W. Pajtler, Jan-Philipp Mallm, Karsten Rippe

## Abstract

**Background:** Spatial transcriptomics (*ST*) technologies are revolutionizing our understanding of intra-tumor heterogeneity and the tumor microenvironment by revealing single-cell molecular profiles within their spatial tissue context. The rapid evolution of *ST* methods, each with unique features, presents a challenge in selecting the most appropriate technology for specific research objectives. Here, we compare four imaging-based *ST* methods – RNAscope HiPlex, Molecular Cartography, MERFISH/Merscope, and Xenium – together with sequencing-based *ST* (Visium). These technologies were used to study cryosections of medulloblastoma with extensive nodularity (MBEN), a tumor chosen for its distinct microanatomical features.

**Results:** Our analysis reveals that automated imaging-based *ST* methods are well suited to delineating the intricate MBEN microanatomy, capturing cell-type-specific transcriptome profiles. We devise approaches to compare the sensitivity and specificity of the different methods together with their unique attributes to guide method selection based on the research aim. Furthermore, we demonstrate how reimaging of slides after the *ST* analysis can markedly improve cell segmentation accuracy and integrate additional transcript and protein readouts to expand the analytical possibilities and depth of insights.

**Conclusions:** This study highlights key distinctions between various *ST* technologies and provides a set of parameters for evaluating their performance. Our findings aid in the informed choice of *ST* methods and delineate approaches for enhancing the resolution and breadth of spatial transcriptomic analyses, thereby contributing to advancing *ST* applications in solid tumor research.

## Background

Single cell RNA sequencing of dissociated cells (scRNA-seq) or nuclei (snRNA-seq) has become a standard method in cancer research to dissect deregulated transcriptional programs as well as cell types and cell fate trajectories [1]. However, in the sc/snRNA-seq analysis spatial relations between cells in their native tissue context are lost. A variety of emerging spatial transcriptomics (*ST*) approaches that acquire molecular gene expression profiles of cells *in situ* reveal the spatial relations between individual cells [2–5]. ST technologies provide novel insight into tumor heterogeneity as well as interactions of tumor cells with their microenvironment [6, 7]. They can be broadly classified into sequencing (*sST*) and imaging (*iST*) based methods. The *sST* analysis employs a readout by sequencing after transcripts have been released from the sample and are captured directly or via hybridized probes, which can be conducted in an unbiased way for the whole transcriptome. The *iST* methods apply multiplex single molecule RNA fluorescence in situ hybridization (smRNA-FISH) approaches in a targeted manner as defined by the probe panel together with transcript identification by imaging. For *ST* experiments of tumor samples, the different methods have their own specific strengths and weaknesses and numerous questions about the best technical implementation of *ST* technologies and the experimental design exist. On the one hand, organism, tissue type as well as sample processing, e.g., formalin-fixed, paraffin-embedded (FFPE) or fresh frozen tissue will affect the results obtained with a given method. On the other hand, there is currently no consensus on how to determine relevant parameters for quality control including the following: (i) The *sensitivity* of the method, which is given by the probability that a given transcript is detected. (ii) The target *specificity* as reflected by the false discovery rate (FDR). (iii) The specific genes and their total number that are covered well in the experiment. (iv) The assignment of transcripts to a cell.

The resolution of the transcriptome analysis and the cell type annotation will depend on the experimental raw data as well as their preprocessing and downstream data analysis. For example, a crucial step in the workflow is the segmentation of cells for transcript assignment and cell type identification. Here, different dyes for staining of nuclei, membrane and whole cells are available but results depend again on organism, tissue type and sample processing. In addition, the microscopy system, e.g., wide-field vs. confocal microscope, used objectives and/or detector sensitivity will affect the quality of the images with respect to the resolution and signal-to-noise ratio and thus segmentation of nuclei and cells. Numerous computational methods such as Cellpose [8], Baysor [9] and Mesmer [10] have been developed for segmentation and their results are highly dependent on the input data.

For high-throughput *iST* commercial instruments with automated imaging and integrated microfluidics or pipetting robotics are advantageous and the following platforms were used in this study: (i) Molecular Cartography (MC) on the MC 1.0 instrument (Resolve Biosicences) [11]. (ii) Multiplexed error-robust fluorescence in situ hybridization (MERFISH) [12] on the Merscope (Vizgen) system [13], which is referred to as “Merscope” in the following. (iii) Hybridization with barcoded padlock probes directly targeting the RNA [14] as implemented on the Xenium Analyzer instrument (10x Genomics) to which we referred to as “Xenium” [15]. A number of reports compared the performance of these and other instruments for FFPE cancer tissue samples [16, 17] as well as mouse brain FFPE [18] and fresh frozen [19] tissue sections. However, a study using the same fresh frozen cancer samples on the different *iST* platforms is lacking. Fresh frozen samples can be advantageous with respect to RNA integrity as well as conducting unbiased single nuclei transcriptome and/or open chromatin profiling by scATAC-seq from the same sample. Here, we applied a comparative *ST* analysis for a case study focusing on medulloblastoma tumors with extensive nodularity (MBEN) [20]. MBENs are a histopathologically defined subtype of medulloblastoma, which is among the most common embryonal central nervous system tumor in children [21, 22]. Due to mutations in the sonic hedgehog pathway MBEN mimics the development of cerebellar granule neuronal precursors and thus features the complete developmental trajectory [20]. This is reflected in the MBEN tissue structure, which is characterized by a segregation into an internodular (proliferating cerebellar granular neuronal precursor-like malignant cells together with stromal, vascular, and immune cells) and nodular compartment (postmitotic, neuronally differentiated malignant cells). In the present study, we conducted an *iST* analysis of the same MBEN patient samples by MC, Merscope and Xenium in comparison to RNAscope HiPlex [23, 24] as a well-established reference for low-throughput *iST*. In addition, snRNA-seq and *sST* on the Visium platform (10x Genomics) were included as methods for an unbiased transcriptome analysis. Based on our experience with these six different methods we identified informative QC parameters and metrics to assess sensitivity and specificity of the different methods. Furthermore, we show how technological differences affect the results and provide guidance for the experimental design for the analysis of fresh frozen tumor samples by *iST*.

## Results

### ST of MBEN samples

The analysis of MBEN samples by different *ST* methods was conducted with fresh frozen tissue from four different patients (**Supplementary Table S1**) that have been studied previously using sequencing, microdissection and spatial technologies [20]. Here, we dissected the distinct MBEN microanatomy with a different cellular composition of the internodular and the nodular compartment in a comparison of different ST methods (**Fig. 1**).

**Figure 1.**
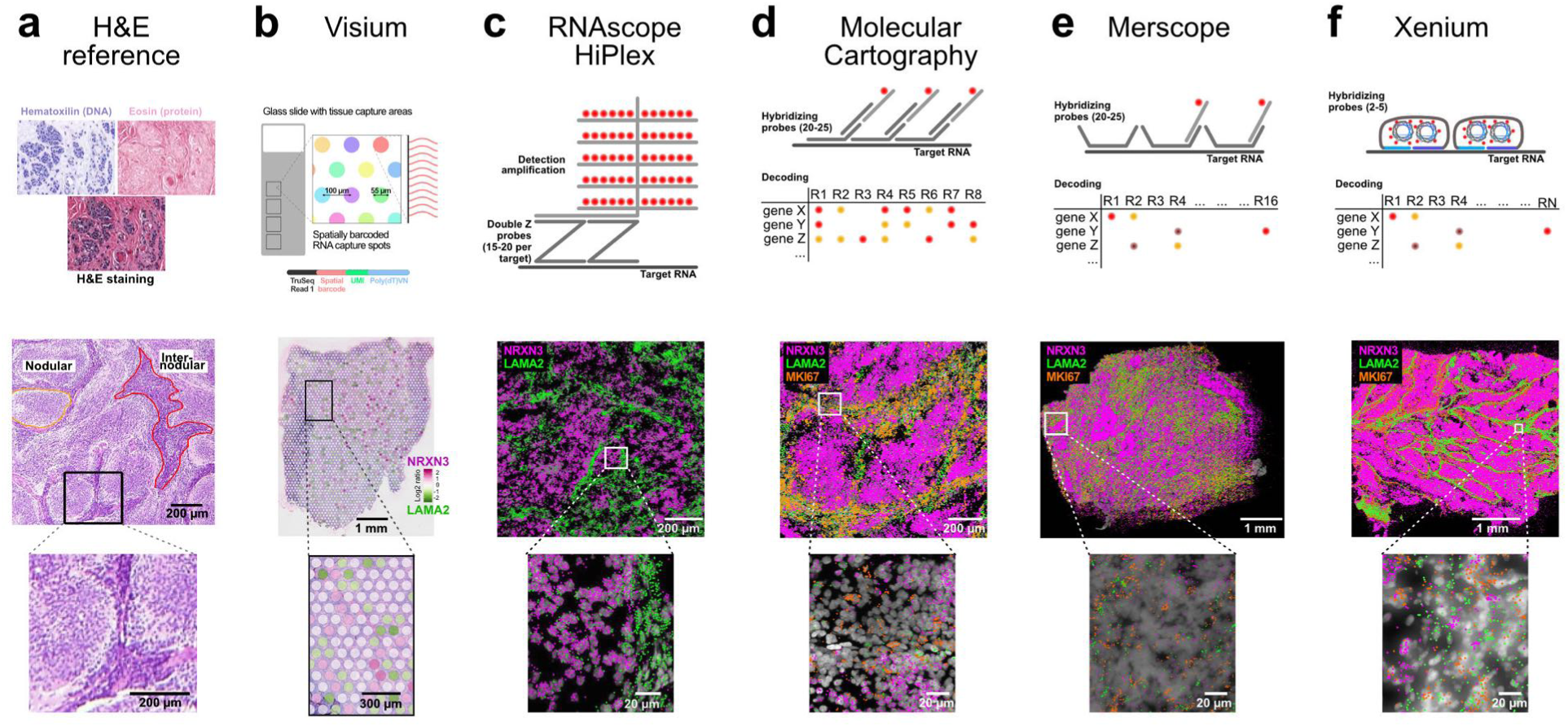
Overview of *ST* technologies compared in this study. Marker genes *NRXN3* (purple color, nodular compartment) and *LAMA2* (green color, internodular compartment) and MKI67 (orange color, proliferating cells) are shown for sample MB295. (**a**) H&E reference staining. (**b**) Visium. (**c**) RNAscope HiPlex. (**d**) Molecular Cartography. (**e**) Merscope. (**f**) Xenium.

Exemplary tissue images are shown for hematoxylin and eosin (H&E) staining (**Fig. 1a**) together with the different *ST* technologies used (**Fig. 1, S1, Supplementary Table S2**). These included Visium (**Fig. 1b**), RNAscope (10 gene panel, **Fig. 1c**), MC (100 gene panel, **Fig. 1d**), Merscope (138 gene panel, **Fig. 1e**) and Xenium (345 gene panel, **Fig. 1f**) (**Supplementary Dataset 1**). All *iST* panels included the 10 genes from RNAscope, and Merscope and Xenium shared 96 genes of the MC panel. The MBEN tumor microanatomy is visible in the H&E staining and was revealed by all *iST* methods on the transcript level by transcription of *NRXN3* and *LAMA2* as marker genes for the nodular and internodular compartments, respectively (**Fig. 1**). The Visium analysis, however, did not provide sufficient spatial resolution to clearly delineate the two different tumor compartments as apparent from the NRXN3/LAMA2 expression ratio (**Fig. 1b**). In addition, we also included snRNA-seq data generated on the Chromium platform in our comparison as a reference for the established and frequently used approach for a single cell transcriptome analysis of solid tumor samples.

### *ST* image acquisition and reimaging of slides

For Visium and RNAscope experiments, the image acquisition is decoupled from the transcript detection and decoding. For these modalities, H&E images were acquired on a slide scanner and the RNA-scope ST data acquisition was conducted by spinning disk confocal microscopy (SDCM).

The commercial MC 1.0, Merscope and Xenium Analyzer instruments provide automated image acquisition on a built-in wide-field fluorescence microscope with some differences concerning objective and camera as well as the software provided for preprocessing. These systems implement different smRNA-FISH protocols, which has some implications for practical usage. Placement of fresh frozen cryosections is relatively easy for MC and Xenium but can be difficult for Merscope if more samples are to be placed on one slide due to the slide architecture (**Fig. S1**). In the MC system readout probes are removed and the whole workflow can be started again in case of a power cut or system malfunction. This is not possible for Merscope, as the fluorophores are bleached. After the ST run, it can be advantageous to conduct a reimaging step to either acquire higher resolution images (e.g., improved DAPI images for cell segmentation, see below) or to include additional image modalities such as H&E, membrane or immunostainings. For MC and Xenium, this is straightforward since the tissue integrity is maintained during the runs and standard slide formats are used (**Fig. S1**). We have developed a workflow for MC and Xenium that reimages the slides by SDCM microscope that are then registered on the spatial transcriptomics map obtained with these systems (**Fig. 2a**). We have not tested reimaging of the Merscope slides since the sample clearing step before the run removes all tissue components other than RNA and DNA and imaging is more difficult due to the custom slide format. To overlay the images of the tissue obtained from a different microscope, images are first stitched and then registered to the DAPI images from the iST analysis of the MC or Xenium system. In this manner, the cell’s transcriptome profile can be integrated with additional readouts.

**Figure 2.**
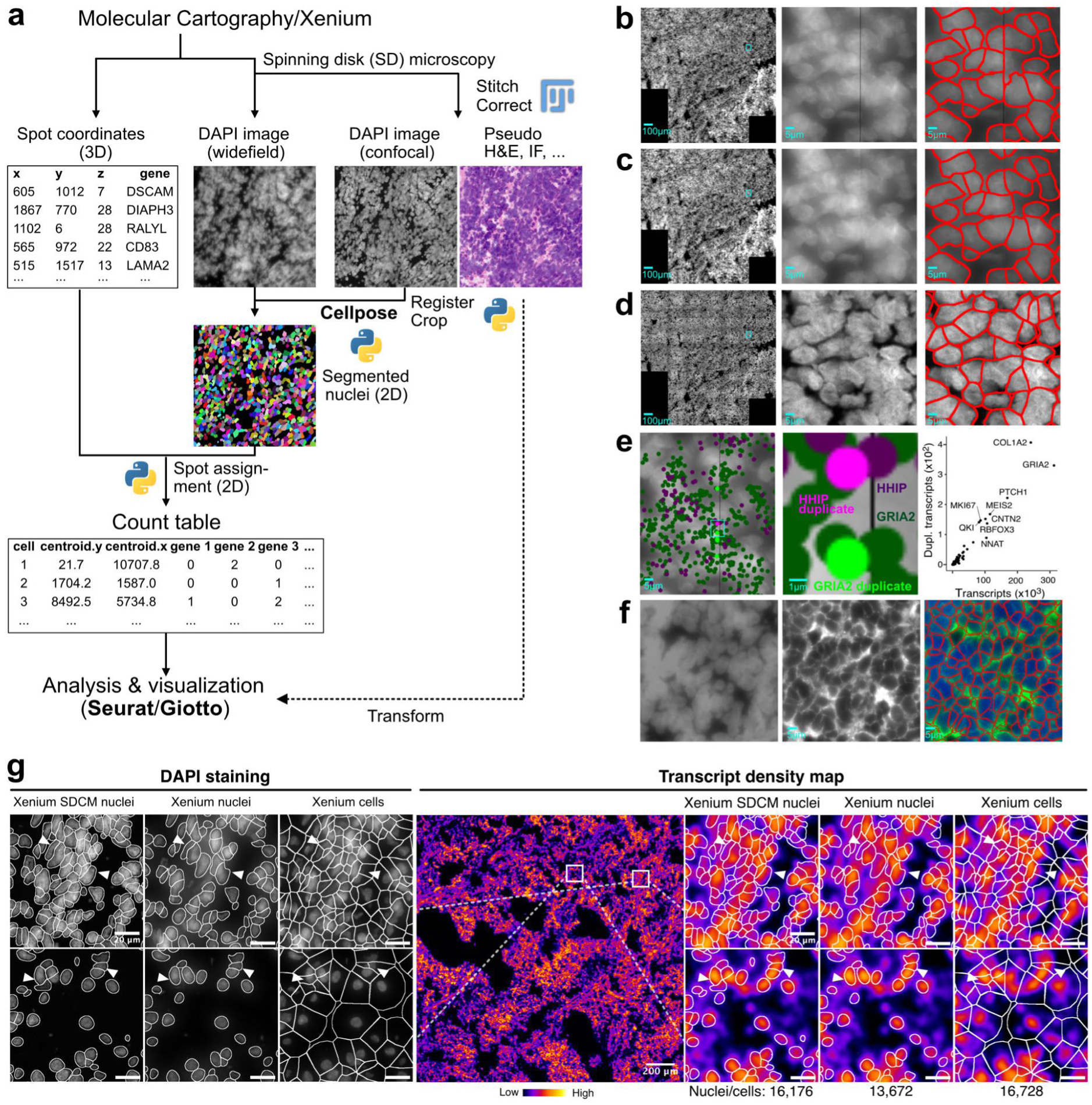
Reimaging and segmentation. (**a**) Reimaging workflow of MC and Xenium slides. (**b**) Widefield overview DAPI image, zoom and segmented image for MC. (**c**) Same as panel **b** but after applying the MindAGap software to fill the non-overlapping line between images. (**d**) Same region as in panel **b** and **c** but after reimaging on the spinning disk microscope. (**e**) The gap between images introduces artifacts for stitching and registration that lead to the artificial generation of duplications for 0.15% of the transcripts. (**f**) Segmentation for Merscope with membrane staining. Left, DAPI stained wide-field image; middle, membrane staining; right, segmentation based on DAPI signal and membrane staining. (**g**) DAPI images (zoom-in of indicated regions) of Xenium slide and segmentation based on SDCM and widefield images for tumor tissue MB266. The region overview is shown as a transcript density. The fraction of transcripts assigned to segmented nuclei or cells were 68% for Xenium SDCM nuclei (Xenium slide reimaged by SDCM), 59% for Xenium nuclei and 95% for Xenium cells. The cell expansion used for the latter segmentation covers almost all transcripts. However, this is associated with artifacts as seen for the cells marked with white triangles. For quantification see **Fig. S2**.

### Image processing and cell segmentation

To assign transcripts to individual cells after segmentation, several image processing steps are conducted. If not noted otherwise, we used segmentation based on DAPI staining (MC 1.0), DAPI and membrane staining (Merscope) and DAPI staining with cell expansion (Xenium analyzer) as default workflows for the different systems. We find that the quality of the DAPI images is crucial with respect to staining, image acquisition parameters (excitation intensity, exposure time, dynamic range of detector) and the image resolution obtained with the wide-field microscopes used in these systems. In particular, it is important to optimize the DAPI signal-to-noise ratio and to avoid a too low signal as well as a fluorescence signal that is out of the dynamic range of the detector. The images provided by the default settings of commercial *iST* systems leave room for improvement in this respect for the very cell dense MBEN tissue sections studied here. To specifically evaluate the effect of image quality, preprocessing and segmentation method, we additionally acquired SDCM images for MC and Xenium (referred to as MC SDCM and Xenium SDCM). In cell dense areas the analysis of the original wide-field images leads to ambiguous results. This is illustrated for the MC workflow in **Fig. 2b-d**. This analysis also revealed stitching artifacts due to non-overlapping images that resulted in black strips splitting cells that span across imaging tiles (**Fig. 2b**), which can be addressed by Gaussian blurring (**Fig. 2c**) but results in partially duplicated cells and transcripts. To further investigate this issue, the confocal images were registered, and transcript duplicates removed (**Fig. 2d**). These duplicates appeared at a low but still detectable frequency of 0.15% across all transcripts (**Fig. 2e**). This type of stitching errors was not detected for Merscope and Xenium systems. Finally, the option provided for Merscope to include a membrane staining in the standard workflow can improve cell segmentation on widefield areas as illustrated in **Fig. 2f**.

In general, segmentation with Cellpose [8] using the DAPI signal yielded good results and the SDCM images provided a 15-30% higher number of segmented nuclei. This is illustrated for the MB266 sample in **Supplementary Fig. S2**. About 71% (MC) and 68% (Xenium) of the overall detected transcripts were located in the segmented nuclei. In contrast, nuclei segmentation on the corresponding widefield microscope yielded ∼10% less assignment of the overall detected transcripts to nuclei (MC, 58%; Xenium, 59%). This can be partially attributed to the lower overall numbers of segmented nuclei on widefield images (∼28% for MC and ∼15% for Xenium in case of MB266). However, it is noteworthy that not only the number of segmented cells or nuclei matters but also their size and shape. The larger the cells the more transcripts were detected. Nevertheless, simple extension of segmented nuclei to include cytoplasmic transcripts resulted in some wrongly assigned transcripts and thus created a mixed transcriptome from different cells (**Fig. 2g**).

### Sensitivity of ST methods

The sensitivity of *ST* methods can be defined as the transcript fraction that is detected. To assess this parameter, we analyzed the distribution of the total number of transcripts detected (“transcripts”) as well as the number of genes (“features”) for the shared 96 gene panel shared between all ST methods except for RNAscope. To exclude the confounding effect of segmentation, we conducted a spatial binning analysis as a segmentation-free approach. Number and type of transcripts were determined within spatial bins (48,74 x 48,74 µm) that correspond to area of a circular Visium spot, which is ∼2,375 µm^2^ in size. The *iST* techniques clearly outperformed the Visium *sST* method with respect to the number of transcripts and features in this comparison (**Fig. 3a**). While increasing the sequencing depth could improve the Visium results, but based on our data, one would still expect that Visium sensitivity will remain well below that of the *iST* techniques. Within the latter group, MC yielded the highest number of transcripts per bin while the number of features was similar for all *iST* methods (**Fig. 3a**).

**Figure 3.**
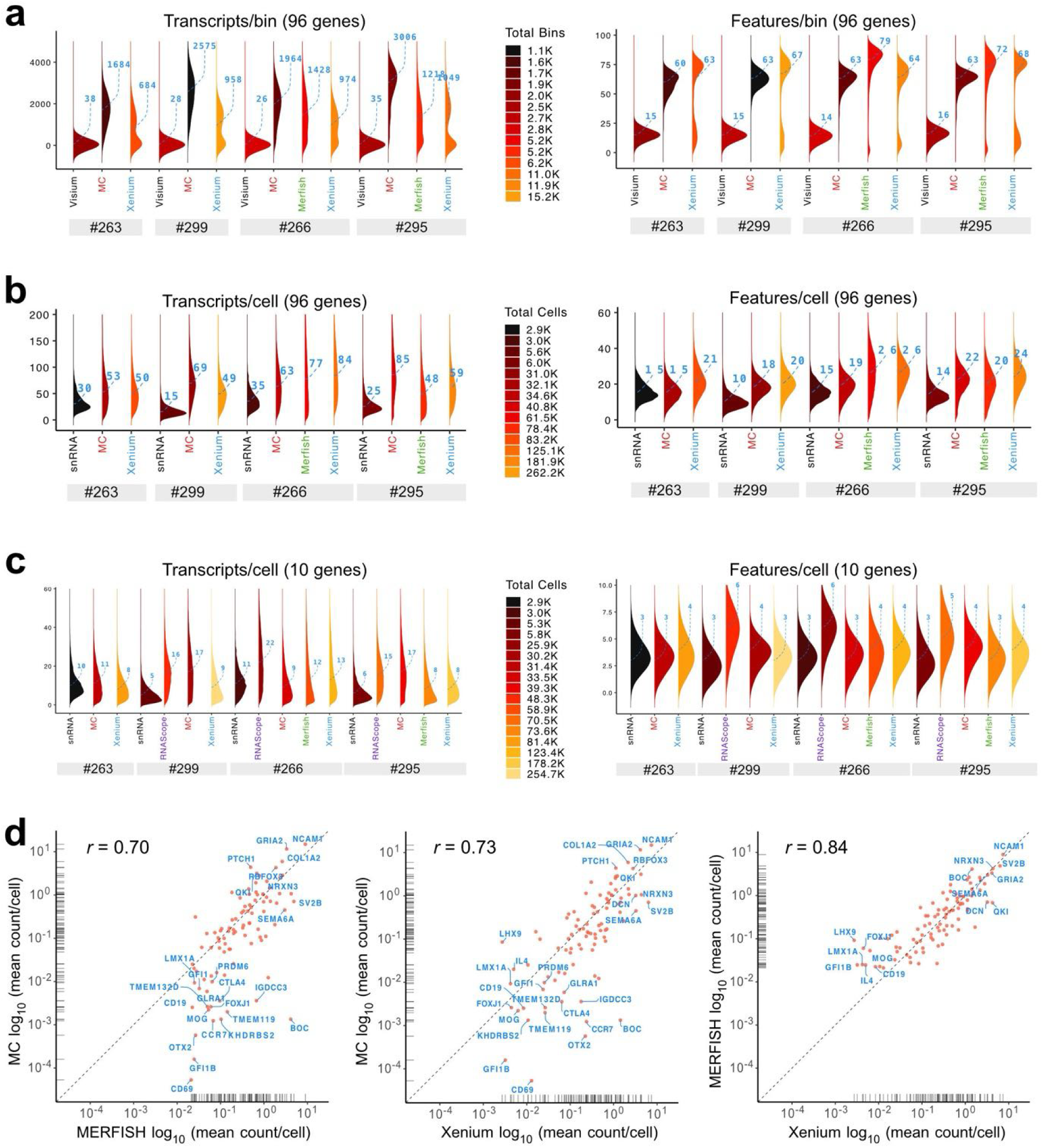
Sensitivity of ST methods. (**a**) Density ridge plots for transcript and feature counts per spatial bin (48.74 µm side length square) being equivalent to the area of one Visium spot. (**b**) Density ridge plots of transcript and feature counts per cell after segmentation for 96 shared genes. (**c**) Same as panel **b** but for the 10 shared genes present in the RNAscope panel. (**d**) Correlations of transcript counts between different *iST* methods. The dashed line depicts the same number of transcripts detected for the two methods compared. Correlations of *iST* with snRNA-seq data are given in **Fig. S3b-d**.

Interestingly, the spatial binning analysis yielded a bimodal distribution for Xenium, which points to the presence of tissue regions with reduced transcript coverage. The cells in the lower transcript distribution appear to be enriched in the outer regions of the tissue (**Fig. S3a**). Next, the number of transcripts or features was computed on a per cell basis for the 96 gene set (**Fig. 3b**). This comparison showed only minor sensitivity differences between the automated *iST* instruments. The well-established RNAscope method consistently yielded high numbers of transcripts/features per cell for the 10 shared genes (**Fig. 3c**). This result confirms its use as a smRNA-FISH reference in the field. We then performed a pairwise comparison of detected mean transcripts per cell between the different *iST* methods for the shared gene set (**Fig. 3d**). For some genes with lower expression levels, e.g., *BOC,* they seemed to be better detected by Merscope and Xenium as compared to MC. The same analysis was applied in reference to snRNA-seq, which has a detection efficiency of 14-15% for the Chromium 3’-RNA v2 chemistry from 10x Genomics used in our experiments (**Fig. S3b-d**). The resulting correlation coefficients of *iST* methods with snRNA-seq were between 0.53-0.63 and somewhat lower than those between the iST methods themselves that ranged from 0.7-0.84 (**Fig. 3d**). On an average, the number of a given transcript per nucleus/cell was 2.3 to 2.5-fold higher for the *iST* methods than for snRNA-seq (**Fig. 3b, S3b-d**). This suggest that the detection efficiency of the *iST* methods is around 33-37%. Overall, the commercial *iST* methods yielded very similar results with MC showing a slightly lower number of counts for less abundant transcripts. When checking the number of transcripts and molecules per cell all *iST* techniques showed higher sensitivity than snRNA-seq as performed with the Chromium v2-chemistry.

### Specificity of iST methods using RNAscope as a reference

To assess *ST* specificity, we used the 10 genes mapped in the RNAscope data as a reference. Correlations of the mean number of transcripts per cell for the 10 genes mapped by RNAscope were calculated. The highest correlation was found between RNAscope and Xenium (**Fig. 4a**). Next, we computed pairwise correlation coefficients between transcripts within a cell for each of the different methods (**Fig. 4b**). According to this correlation analysis, the RNAscope data reflected the MBEN tissue microanatomy described in ref. [20] very well. Marker genes of the nodular compartment (*RBFOX3*, *NRXN3*) as well as those of the internodular compartment (*GLI1*, *TRPM3*, *LAMA2* and *PTCH1*) showed high positive correlations among each other but were not correlated or anti-correlated between these two groups. The Merscope data were most similar to the pattern of (anti-)correlations between gene pairs seen in the RNAscope data (coefficient of determination R^2^ = 0.72), while it was somewhat less apparent for the other methods (MC, R^2^ = 0.45; Xenium, R^2^ = 0.42) (**Supplementary Table S3**). This type of assessment is based on prior knowledge about the spatial expression patterns of a given tissue. It can be implemented after cell segmentation as a quality assessment for specific marker genes that display distinct spatial relations as demonstrated here for MBEN.

**Figure 4.**
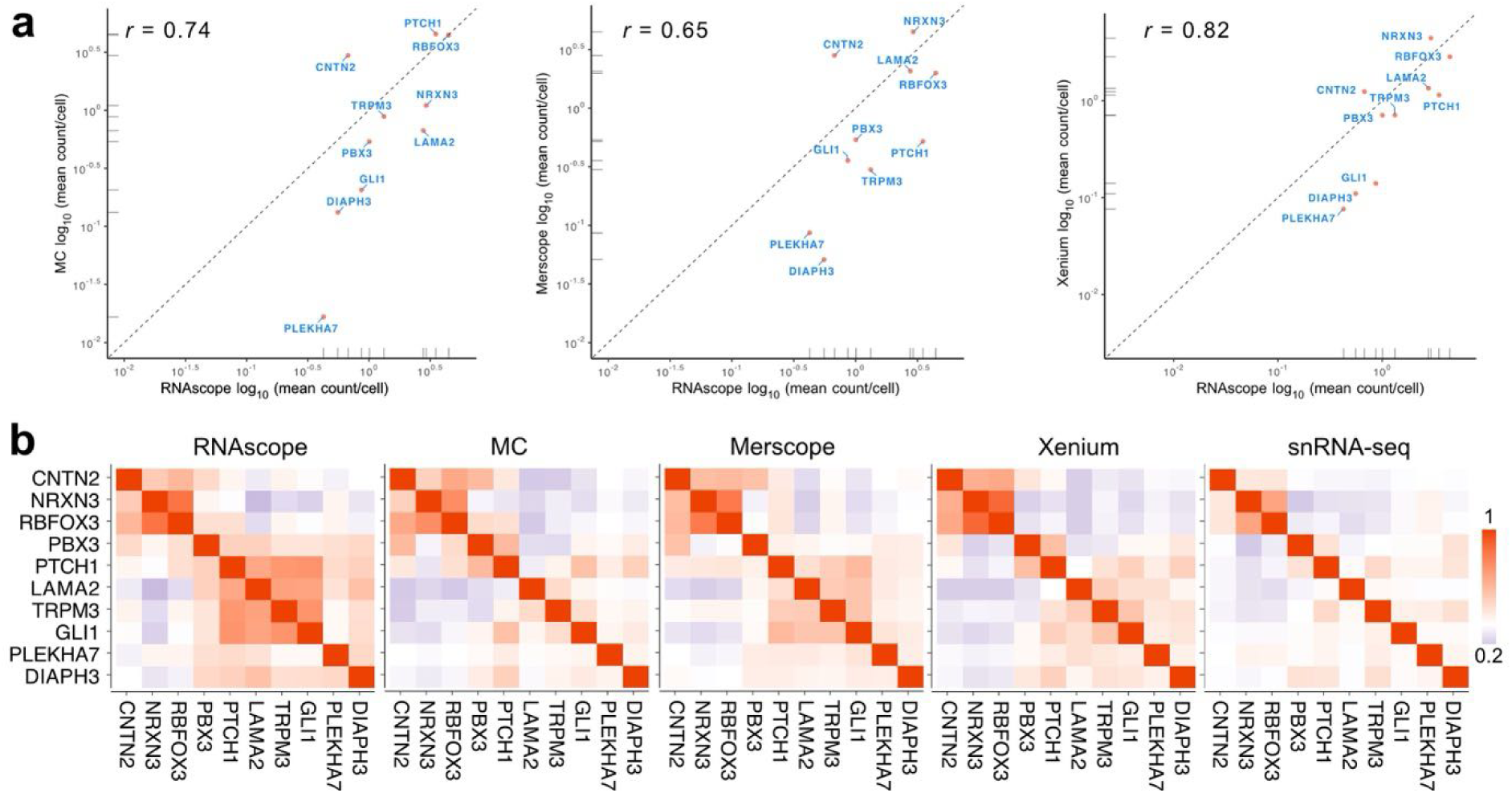
Comparison with RNAscope. (**a**) Correlation of gene expression for the different *iST* methods with RNAscope for the 10 shared genes. The dashed line depicts the same number of transcripts detected for the two methods compared and shows that 7/10 (MC) and 8/10 genes (Merscope, Xenium) had a higher number of transcripts per cell/nucleus while CTN2 was detected better with all *iST* methods. (**b**) Analysis of marker gene co-expression from the pairwise correlation (Pearson) coefficient.

### Specificity of iST methods inferred from background probes

Next, we assessed specificity by relating the signal obtained from fluorescently labeled control probes referred to here as background probes that lack a complementary sequence in the sample to the target genes of the panel on different length scales as depicted in **Fig. 5a**. It is noted that the background probes were those provided by the manufacturers and information on their sequences is lacking. In addition, the three *iST* methods employ different controls (**Supplementary Table S4**). MC and Merscope depend on binding of numerous probes to gain sufficient signal. Thus, false positive signals usually occur via the read-out probes rather than the primary probes. For Xenium, due to the amplification of the signal from a single probe, both off-target binding of primary and secondary probes needs to be considered. Accordingly, also unspecific primary probes are included in the kit.

**Figure 5.**
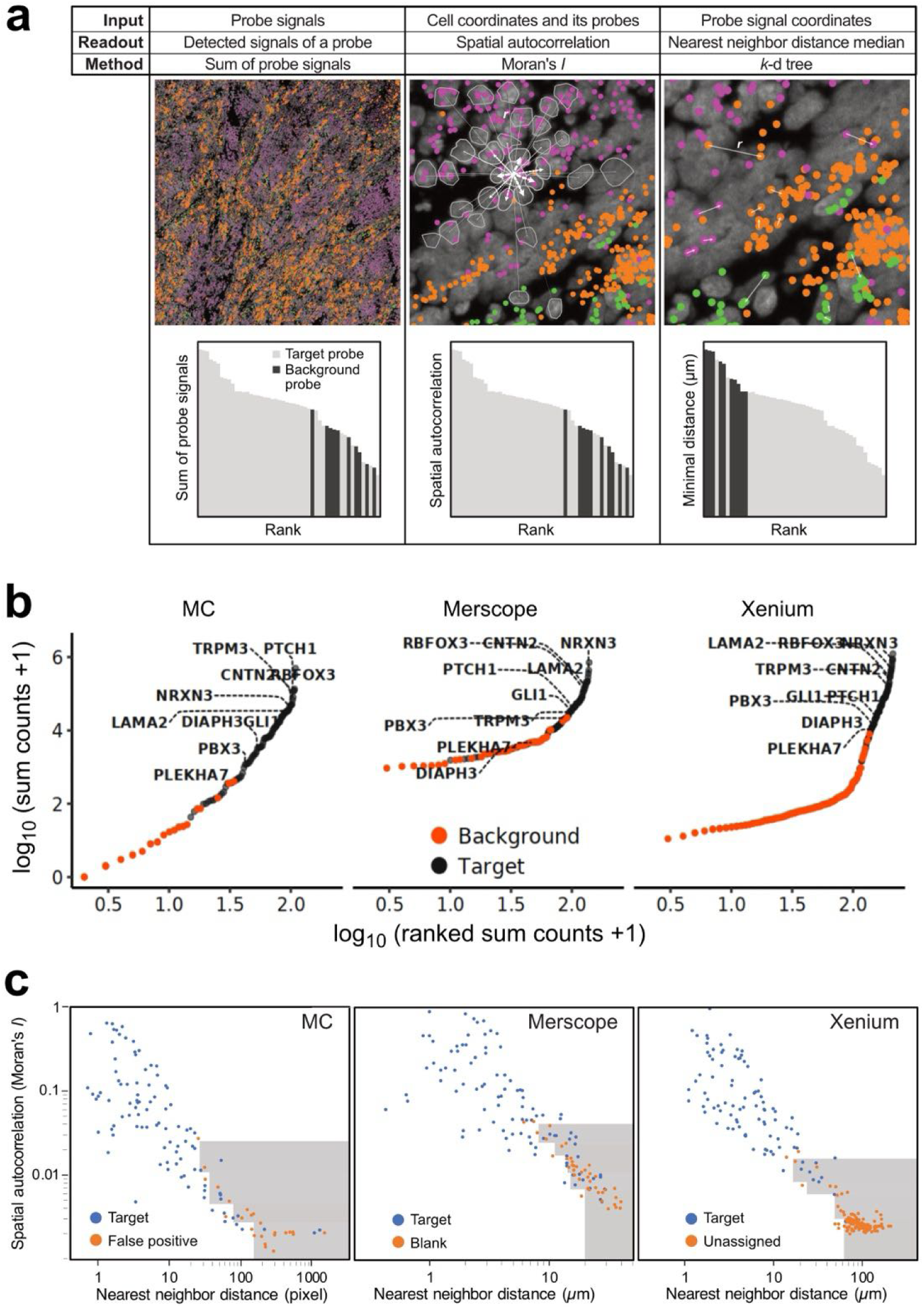
Specificity analysis from a comparison of target and background probes. Data are shown for MBEN 266. (**a**) Specificity analysis on different length scales. Coordinates for three different transcripts are depicted in purple (*NRXN3*), green (*LAMA2*) and orange (*MKI67*) color. *Left*: whole tissue analysis where target and background probes are ranked after summing up all signals detected for a given probes. *Middle*: spatial autocorrelation of probe signal computed using Moran’s *I* at the cell level. This value increases if a given cell’s signal (marked by outgoing distance vectors) is similar to the average value of neighboring cells at distance *r* as indicated by the connecting vectors, which is weighted with 1/*r* between two cells (vector thickness indicates higher weights). *Right*: minimal distance to the next probe signal of the same type. This parameter can be used to identify clusters at subcellular resolutions originating present in rare and isolated cell types. Exemplary pairs of transcripts are marked with white arrows. (**b**) Distribution of target and background signal plotted against the signal count. See **Fig. S5** for additional datasets. (**c**) Analysis of the spatial distribution of target and background probes. The spatial autocorrelation (Moran’s *I*, scaled to a range from 0 to 1) was plotted against the median nearest neighbor distance. Higher values of Moran’s *I* and lower values of the nearest neighbor distance are indicative of a non-random distribution. Grey area indicates low confidence probes based on 0.05 percentile of the nearest neighbor distance in a given range of Moran’s *I*.

By comparing the sum of all signals for a given probe across the whole tissue, we identified 29±8 (MC), 43±2 (Merscope) and 18±2 (Xenium) probes for which the signal range was within the range of signals obtained with the background probes (**Fig. 5b, S4a**). Of these, the genes *GFI1*, *LMX1A*, *IL4*, *FOXJ1*, *CD19*, *TMEM119*, *MOG*, *CD69* and *GFI1B* displayed a consistently low expression value for all three iST technologies, which could point to true negative signals. Based on the calls of target and background probes we computed global, segmentation-free FDR values of 0.41±0.2 % (MC), 5.23±0.9 % (Merscope) and 0.47±0.1 % (Xenium). According to these averaged global FDR estimates, specificity is very similar for MC and Xenium with a higher FDR value determined for Merscope.

The averaged FDR value does not account for the presence of specific signals that are simply lowly abundant. Accordingly, we evaluated the spatial distribution of the target probes by computing their spatial autocorrelation using Moran’s *I* [25] as well as the minimal distance between probe signals against the same target (**Fig. 5a**). This analysis was conducted with the rational that a false positive signal due to technical issues would be randomly distributed (*I* = 0) while a lowly abundant true positive signal (e.g., a lowly expressed marker for a niche cell type) would show some enrichment (I > 0) and/or display clustering at the subcellular level within isolated rare cell types that would yield a low minimal distance. Thus, by combining spatial autocorrelation signal and nearest neighbor distances specific cut-offs can be used to identify targets that reflect a lowly abundant specific signal that is not randomly distributed in tissue space (**Fig. 5c, S4b**). The spatial autocorrelation is conducted at the resolution of individual cells and their neighboring cells, whereas the distance of a transcript to its next nearest neighbor covers also subcellular distances. This distance would be small for transcripts present mostly only in isolated rare cell types that are scattered across the tissue sections. We used a 0.95 percentile cut-off for a given Moran’s *I* range (four distinct ranges for each technique based on **Supplementary Dataset 2**). The number of confident transcripts increased for all techniques as compared to the expression level analysis. Around 7, 12 and 17 transcripts failed the threshold for Xenium, MC and Merscope respectively. This points to a slightly noisier signal for the latter method, which is in line with its higher average FDR value.

### Detection of cell types across platforms

To compare the six different methods with respect to cell type identification we followed standard clustering workflows, assigned cell types based on the expression signatures identified in our previous MBEN study [20] and visualized the data as UMAPs (**Fig. 6a-c, Fig. S5**). The overall cell type annotation was very similar for the *iST* methods and the same major cell types were found across all platforms. Inspection of the cluster heatmaps, however, revealed some difference in the detection efficiency of single transcripts that affect the cell type assignment (**Fig. S5d-f, Supplementary Dataset 3**). For example, the *TULP1*, *KHDRBS2* and *CD19* were only detected in MC, Merscope and Xenium respectively. Accordingly, cell type annotations differ between the technologies mainly due to transcripts that are detected on one platform but not on the other as for example for CD19 and B cells. Subclustering of the “differentiated neuronal-like” annotation occurred for both Merscope and Xenium, while the stromal compartment was subclustered in MC. Whether these subclusters indeed represent distinct cell types/states is would require further investigations. The cell type annotation and corresponding coloring was also used for visualization of their spatial distribution on the images and exemplary regions are shown in **Fig. 6d-f**.

**Figure 6.**
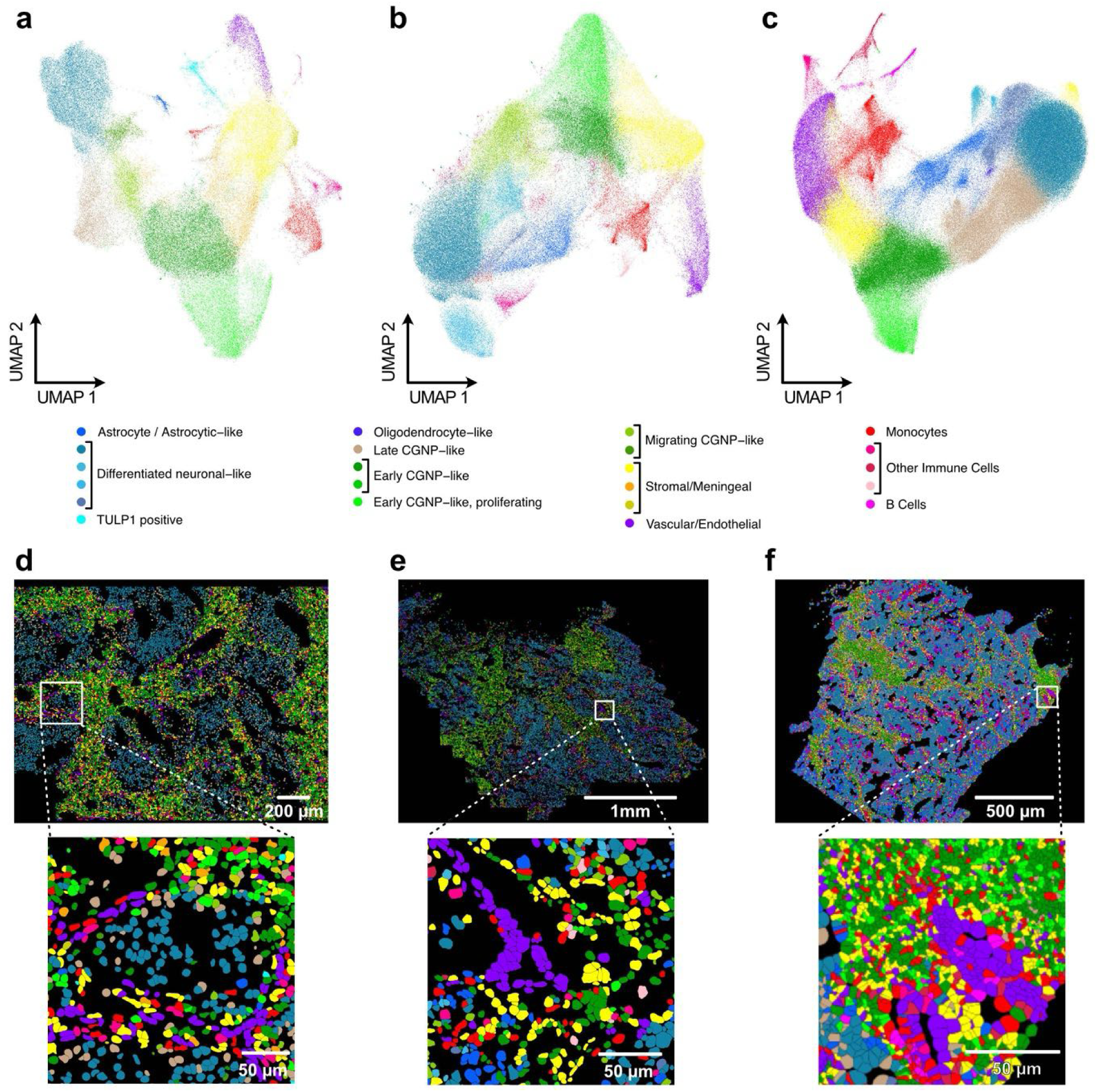
Clustering and cell type annotation for iST methods. Clustering was based on the shared set of 96 genes in samples MB266 and MB295 and MB266. The joint cell type annotation was based on the expression signatures of the different clusters that is described in **Supplementary Fig. S5**. (**a**) Clustering and UMAP visualization for Molecular Cartography. (**b**) Same as panel **a** but for Merscope. (**c**) Same as panel **a** but for Xenium. (**d**) Images with cell coloring according to cluster. (**e**) Same as panel **d** but for Merscope. (**f**) Same as panel d but for Xenium.

### Implementation of additional readouts after iST analysis

While the *ST* analysis provides a wealth of information on molecular cellular profiles in their spatial tissue context, the corresponding studies typically require the integration with other readouts. To accomplish this, a frequently used approach is to prepare consecutive tissue sections used for *ST* and other readouts. However, in many instances, the cell-by-cell assignment of the consecutive sections is cumbersome and works only in some areas. Alternative approaches involve performing additional readouts on the same tissue by either reimaging and subsequent image registration (MC and Xenium) or including additional custom readouts directly in the *ST* run (Merscope). This is described here for three examples.

The first one involves a virtual H&E staining of tissue after the MC run (**Fig. 7a**). Conventional H&E staining after the *iST* run is compatible with both MC and Xenium, however it will prohibit subsequent acquisition of additional fluorescence signals since the broad fluorescence emission of hematoxylin interferes with additional signal. This can be circumvented by fluorescence imaging of DNA via DAPI staining (λ_ex_ = 405 nm, λ_em_ = 421±23 nm) and eosin (λ_ex_ = 488 nm, λ_em_ = 521±19 nm) followed by a transformation of these signals into a virtual H&E staining (**Fig. 7a**) [26, 27]. However, contributions like cell type specific shades of pink seen with eosin in brightfield images or differences in cell autofluorescence cannot be fully accounted for with the approach used here.

**Figure 7.**
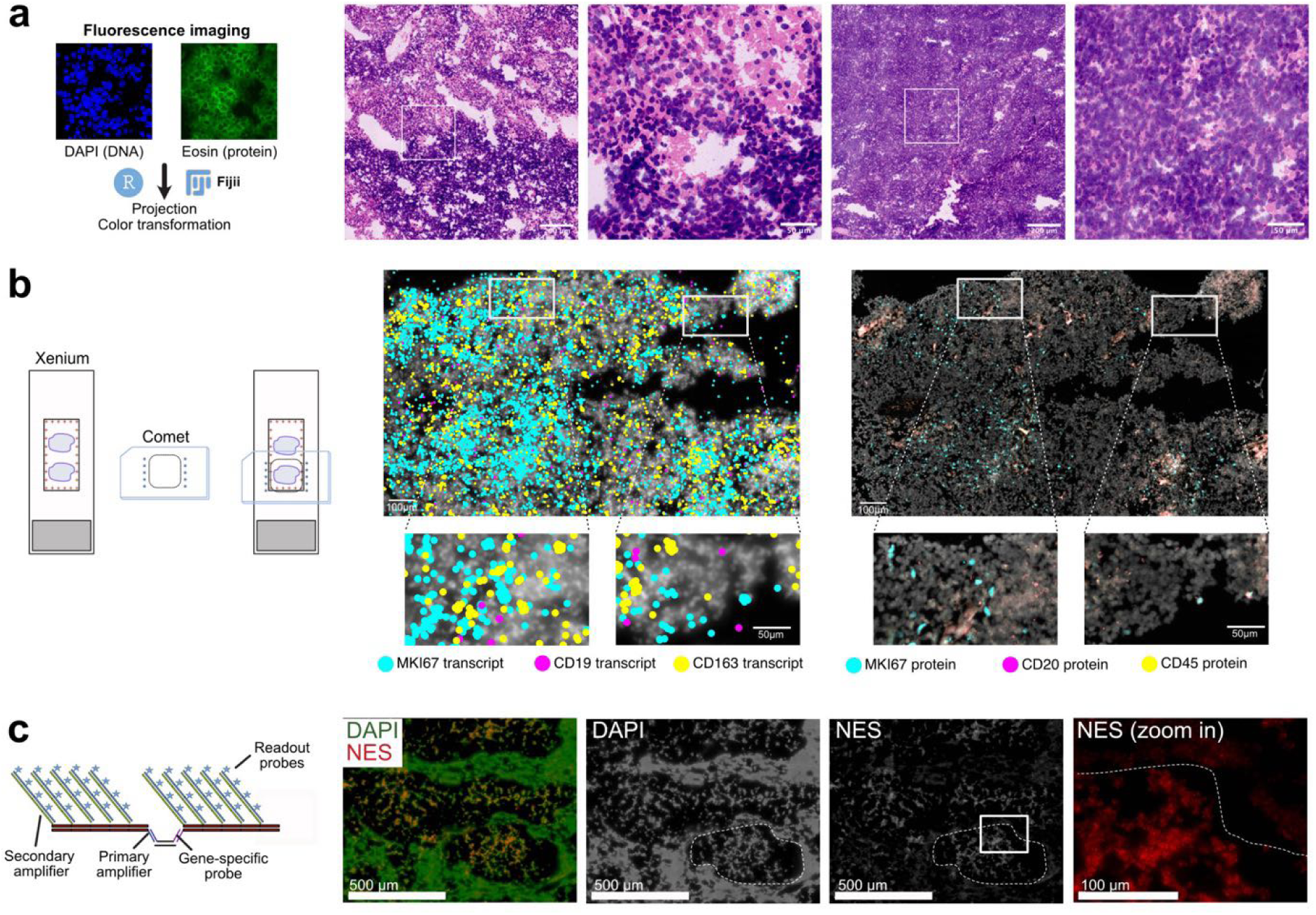
Additional readouts after iST analysis. (**a**) Virtual H&E staining of MBEN tissue after MC run. (**b**) Immunostaining on the Comet system from after the Xenium run. Transcript signal is given as a single dot while the protein image reflects the original fluorescent signal. (**c**) Amplified readout of nestin (NES) as an exemplary custom RNA via the auxiliary channel on the Merscope system.

In **Fig. 7b** we show that the Xenium slide can also be used for subsequent multiplexed immunostaining on the Comet platform (Lunaphore) (**Supplementary Table S5**). As a test case we used the validation of the *CD19* transcription signal, which was detected on the Xenium but not on the MC and Merscope system. By using immune and B-cell specific antibodies as well as Ki67 to stain cycling cells with the Comet system after a Xenium run, the immunostaining confirms the presence of CD19+/CD20+ positive B cells both on the transcript and protein level.

The third example is the detection of a custom gene with the Merscope using signal amplification [28] (**Fig. 7c**). The primary probe carries overhangs to which a primary amplification probe hybridizes that in turn provides a binding platform for a secondary amplifier (**Supplementary Table S6**). The secondary amplifier can be detected with auxiliary probes in the Merscope chemistry. Nestin (NES) as an exemplary custom RNA target was detected via an auxiliary probe on the Merscope system and showed enrichment in the nodular structure (dashed outline). Signal amplification enabled the detection of NES by using only two primary probes as opposed to 30 in the original workflow. This workflow could be used to detect for example short transcripts or gene fusions.

## Discussion

Our comparative analysis of six different *ST* methods used MBEN cryosections as a case study. Because of its characteristic microanatomy with two distinct tumor cell compartments, this entity is particularly interesting for an *ST* analysis of the interplay of proliferation, migration and differentiation of cancer cells [20]. In addition, an *ST* analysis revealed important information for the spatial relation of tumor subclones and proliferating tumor cells in Group 3/4 medulloblastomas that have been reported in two recent studies [29, 30]. Our present work provides valuable insights for the application of *ST* technologies specifically to tumor cryosections, which can differ from non-malignant tissue, as for example by a high local cell density seen here for MBEN. Other tissue and sample types like mouse brain sections or FFPE samples have different requirements for optimizing the *ST* analysis. Furthermore, despite our efforts to standardize protocols and maximize comparability, technical variability is introduced from differences in the complex experimental and data analysis workflow. Factors like tissue handling, staining efficiency, and the calibration of the instrument imaging settings impact on sensitivity and specificity. Accordingly, they might influence the results in an experiment dependent manner that is not reflecting platform related differences. Finally, *ST* technologies are rapidly evolving and undergo constant improvements of chemistry, experimental data acquisition and updates of the instrument specific software. For example, the MC 1.0 instrument will be replaced by a new system and new chemistry versions were released continuously for multiple workflows. Thus, our study is not suited to select “the best” technology. Rather, we see its value in identifying current key differences between the methods together with critical steps in the workflow that warrant consideration for the experimental design as well as in the assessment of the results. To guide selecting a method for a specific application we have summarized selected features of the different platforms in **Table 1**.

**Table 1.** Feature overview of commercial *iST* platforms.

| Control | MC | Merscope | Xenium |
| --- | --- | --- | --- |
| Detected targets per cell <sup>a</sup> | 21±2 | 23±4 | 25±1 |
| Transcripts/cell <sup>a</sup> | 74±11 | 62±14 | 71±13 |
| Correlation with RNAscope <sup>b</sup> | Medium | High | Medium |
| Targets counts overlapping with background signal <sup>a,c</sup> | 29±8 | 43±2 | 18±2 |
| Average FDR (%) | 0.41±0.2 | 5.23±0.9 | 0.47±0.1 |
| Probes with low specificity <sup>d</sup> | 12±3 | 17±3 | 7±3 |
| Re-imaging | Yes <sup>e</sup> | No | Yes |
| Time to data (days) | 4 | 2-3 | 2 |

<sup>a</sup> Data are averages of median values and their standard errors and refer to the shared 96 gene panel.
<sup>b</sup> Based on 10 gene panel shared with RNAscope.
<sup>c</sup> Different background probes were used for the different technologies (**Table S4**).
<sup>d</sup> Probes that displayed a signal intensity in the range of the background value, had low spatial autocorrelation and a higher minimal distance to its nearest neighbor as described in the context of **Fig. 5**.
<sup>e</sup> Slide is glued to chamber.

In our experience, MC is frequently used to study a limited number of samples with its standard reagent kit for one run (8 tissue sections on one slide) and a panel of 100 custom genes. This format is quite flexible, and makes it well suited for validation experiments. Vizgen allows for larger customized panels of up to 960 genes, while many Xenium users aim to acquire large scale data sets with respect to the number of samples/tissue area analyzed. From a practical point of view, we experienced easy and user-friendly protocols for both MC and Xenium with simple washing and incubation steps. Merscope requires not only more time but also optimization for clearance, quenching and stable gel formation. The latter can result in the loss of the samples in case of gel detachment. The systems (microscope and liquid management) usually worked robustly. However, issues occurred with sample transport and liquid handling for MC due to the modular setup with a robotic arm and liquid pipetting robot. Alignment of the objective for the Merscope required attention and re-adjustment while the Xenium displayed malfunctions due to liquid leakage. Overall, in our hands, the least aborted runs were observed for the Xenium system. However, as technology and associated costs are rapidly evolving these scenarios are only snapshots of the current state. Accordingly, we focus here on identifying guidelines for the experimental design and analysis based on our case study.

### Including snRNA-seq data

The snRNA-seq data are a very good reference to select the probe panel for the targeted *iST* methods. The *sST* methods such as Visium, Slide-seq/Curio Seeker [31] and others could also provide this information. In our hands, the snRNA-seq approach is more straightforward since it can use established single cell sequencing workflows with the caveat that data will depend on the quality and amount of the isolated nuclei used as input. New tools for panel design have been released to adjust for platform specifications [32]. Additionally, snRNA-seq data are also a very good reference to assess the quality and coverage of the *ST* data.

### Number of probes

With good *a priori* information (e.g., from snRNA-seq) even relatively small probe panels like the 10 genes used for RNAscope already resolved the main cell types (**Fig. S5b**). Thus, 100 well selected genes might be more informative than a several fold higher number of probes from a more generic catalog panel. Again, the snRNA-seq data provide an excellent reference to test the suitability of the *iST* probe panel. The latter can be used to conduct a probe specific subsetting, clustering and UMAP visualization from the snRNA-seq data and then evaluate the quality of the resulting cell type resolution [33].

### Cell segmentation

In comparative *ST* studies mouse brain tissue is frequently used as a reference. However, cancer tissue like the MBEN sections studied here typically have a much higher cell density, which makes segmentation more challenging. We find that optimizing the DAPI imaging, both with respect to staining and image acquisition, can largely improve results. Furthermore, segmentation of nuclei with Cellpose yields robust results and, due to the high cell density of the tissue, the loss of information from transcripts outside of nuclei, had no apparent negative effect on the downstream analysis in our study. In addition to Cellpose, other powerful cell and nuclei segmentation tools are available. These include Mesmer [10] and Stardist [34] as well as *iST* specific tools like Baysor [9], SCS [35] and BIDCell [36]. We also recommend testing whether optimizations on the cell segmentation part actually translate in improvements of the results of interest in the downstream analysis since this part of the workflow can become very time consuming. Finally, several pipelines have been published which enable the automated preprocessing of *iST* data like Molkart for MC [37] and the technology agnostic SOPA [38].

### Sensitivity

*T*he unbiased *sST* analysis by Visium covers the complete transcriptome. However, its detection sensitivity and spatial resolution were insufficient to resolve the MBEN microanatomy, which largely limits its application in our case study. Other sequencing-based methods like Seq-Scope [39], Stereo-seq [40], Slide-seq/Curio Seeker [31] and Visium HD provide higher subcellular spatial resolution. However, it is noted that sufficient RNA molecules need to be captured per area, which might require spatial binning at the expense of spatial resolution. Furthermore, sensitivity of sequencing based spatial technologies depends on the read depth, in contrast to the imaging-based workflows that always have the full coverage in terms of transcript number. Improvements in image resolution, e.g., by structured illumination microscopy (SIM) [41] or other super-resolution techniques might overcome crowding effects that can limit sensitivity and/or specificity.

In general, the sensitivity of all three commercial *iST* platforms used in our study was high and very similar. For the non-amplified MERFISH method values of 50% [42] or 80% [43] have been reported previously for the detection rate with cell line samples using a custom microscope and expansion. Using snRNA-seq as a reference for our analysis of transcripts per cell (**Fig. 3b, S3b-d**), we estimated that the averaged detection efficiency in our experiments was between 33-37 %. It is noted that the sensitivity depends on the integrity of the RNA, which is usually very good for fresh frozen material. In contrast, RNA degradation can be very significant in FFPE samples, which is likely to translate in significant differences between technologies as reported recently [17].

### Specificity

The specificity is dependent on both probe and tissue features and thus difficult to assess in a quantitative manner. The background probes included in runs with the different systems typically show some overlap with low signal target probes. Thus, there is no apparent cutoff in terms of true negatives and false positives with respect to the signal intensity. Including the spatial distribution of probes as an additional parameter to distinguish between random and more localized and bona fide specific binding is helpful but also requires a probe-by-probe interpretation of the results. As shown here, the assessment of the distribution patterns using spatial autocorrelation and next nearest neighbor distance analysis can provide valuable insight in probe specificity irrespective of the global expression level which can give misleading results.

### Reimaging and inclusion of additional readouts

Reimaging of the tissue to improve the image quality for segmentation and/or to include additional readouts benefits from the non-destructive sample processing and slide format of MC and Xenium. The method of choice for yielding images with improved resolution and signal-to-noise ratio are SDCM systems with highly sensitive sCMOS or EMCCD cameras. Imaging with point confocal microscope was found to be too slow for larger tissue areas in our hands. Alternatively, it is also possible to apply an additional analysis on automated commercial widefield platforms as shown here for the immunostaining on the Comet system after the Xenium run **Fig. 7b**. The Merscope samples are less suited for reimaging due to their slide format and clearing of the sample. However, the platform integrates membrane staining in the standard workflow (a feature expected also to become available for the other systems) and provides additional custom readouts via its auxiliary channels. The latter can be used flexibly for custom readouts as for example with signal amplification as illustrated in **Fig. 7c** or antibody staining. Currently, in addition to ST methods, different spatially resolved (epi)genome, proteome and metabolome readouts are becoming available that are in many instances non-destructive and compatible with each other [4, 7, 44]. Accordingly, both for commercial instruments as well as for custom academic workflows, spatial multi-omics approaches are emerging that will further increase the depth of insight that can be gained from the analysis of the same tissue section as opposed to combining the separate analysis of consecutive sections.

In summary, we find that for cryosections of tumor tissue, all three *iST* methods performed very well in terms of their sensitivity and specificity in our case study. In addition, the spatial distribution of cell types annotated based on the shared set of 96 genes studied yielded very similar pictures of the MBEN tissue microanatomy and cellular neighborhoods. Accordingly, selecting one over the other technology platforms would depend on the other criteria discussed above that arise from differences in the technology and their implementation as well as associated requirements for the practical work.

## Acknowledgements

We thank Lambda Moses for discussions on the spatial autocorrelation analysis. We are grateful for the technical support by the Genomics and Proteomics Core Facility and the Omics IT and Data Management Core Facility of DKFZ. This work was supported by the Health + Life Science Alliance Heidelberg Mannheim within the MULTI-SPACE and Explore!Tech programs (CloneSpace, STARnP and LiverMap) and by the German Federal Ministry of Education and Research (BMBF) via project SATURN3 (01KD2206B) within the National Decade against Cancer program. Data storage service SDS@hd was supported by the Ministry of Science, Research and the Arts Baden-Württemberg and the DFG through grant INST 35/1314-1 and 35/1503-1 FUGG. MB was supported by the Deutsche Forschungsgemeinschaft (DFG, German Research Foundation) within the program NFDI4BIOIMAGE (NFDI 46/1 501864659) of the German National Research Data Infrastructure (NFDI) and DRG by the German Academic Scholarship Foundation (Studienstiftung des Deutschen Volkes) and the Mildred Scheel Doctoral Fellowship program of the German Cancer Aid (Deutsche Krebshilfe).

## Author contributions

Acquisition of data: AR, JPM, SS, PS, DK, SJW. Analysis, and interpretation of data: AH, MB, AR, JPM, SJW, KR, DRG, KO. Design and conceptualization: KR, JPM. Writing of the original draft: KR, JPM. Reviewing & editing of manuscript: all authors. Supervision: KR, JPM, AR, KWP.

## Ethics approval and consent to participate

This study was performed after approval by the ethics committee of the Medical Faculty of Heidelberg University. All experiments in this study involving human tissue or data were conducted in accordance with the Declaration of Helsinki and relevant national and international ethical regulations.

## Availability of data and analysis scripts

An overview of the supplementary data associated with this manuscript is given in **Supplementary Table S7** and **S8**. Primary data and data from the downstream analysis including those from ref. [20] are available from the following locations: snRNA-seq, GEO accession number GSE239854; RNAscope, BioImage Archive accession number S-BIAD826; MC, BioImage Archive (accession number S-BIAD825) and GEO (accession number GSE247736); Additional MC data as well as Merscope and Xenium, BioImage Archive (accession number S-BIAD1093); Visium data and Seurat objects have been deposited from Zenodo at https://doi.org/10.5281/zenodo.10863259. The data analysis software used is listed in **Supplementary Table S9** and **S10**. Custom analysis software tools are available via the Github repositories https://github.com/scOpenLab and https://github.com/RippeLab/MBEN.

## Material and Methods

### Tissue samples

MBEN samples MB263, MB266, MB295 and MB299 used in this study have been described previously [20] and their analysis with the different technologies is listed in **Supplementary Table S1**. Cryosections of 10 µm thickness were acquired with a Cryostar NX50 (Epredia) cryostat at a cutting temperature of -15 °C for all technologies. Subsequent storage and processing was performed following the standard protocols provided for each workflow as described below.

### snRNA-seq

The snRNA-seq data were from ref. [20] (accession number GSE239854) and acquired on the Chromium drop-seq platform using 3’-Single Cell RNA-sequencing v2 kit (10x Genomics).

### Visium

Tissues slices of 10 µm were placed on the Visium slides and fixed with methanol at -20°C. After H&E staining the samples were imaged using an Olympus VS200 scanner and the tissue was lysed for 4 min according to the tissue optimization results that were obtained previously. Visium libraries were generated following the manufactureŕs recommendations. Libraries were quantified using Tapestation and Qubit and sequenced on a NovaSeq 6000 machine pooling four libraries per lane.

### RNAscope HiPlex

The RNAscope HiPlex data involved 10 targets (**Supplementary Dataset 1**) and were acquired as described in ref. [20] using the RNAscope HiPlex assay (ACD/Biotechne) according to the RNAscope HiPlex Assay User Manual (324100-UM) from the manufacturer with minor adaptions. For MB266 and MB299, four transcripts (labeled with Alexa488, Atto550, Atto647 and Alexa750 fluorescent dyes) were imaged in three imaging rounds while for MB295, three transcripts (Alexa488, Atto550, Atto647 and DAPI) were imaged in four imaging rounds using the RNAscope HiPlex Alternate Display Module (R1-R4). Flatfield-correction was conducted based on DAPI images and used for nuclei segmentation with Cellpose as described below. Spot-calling of transcripts was conducted with RS-FISH. Called transcripts from all rounds and colors were reformatted and concatenated in an output file in MC format.

### Molecular Cartography

The MC data were from ref. [20] and were acquired with the probe set given in the **Supplementary Dataset 1**. They were reanalyzed by using the restained with DAPI and reimaged on the Andor Dragonfly SDCM system. Image processing followed the workflow depicted in **Fig. 2a**. It comprised stitching, correction and registration as described in further detail in the image analysis section below. The resulting images were then processed with the resolve-processing pipeline (https://github.com/scOpenLab/resolve_processing). The images were first processed with the contrast limited adaptive histogram equalization (CLAHE) [45] with a kernel size of 127 a clip limit of 0.01 and 256 bins. The resulting images were then segmented using CellPose2 with the “cyto” model, a probability threshold of one and a cell diameter of 70. After cell segmentation the transcripts were deduplicated with the MindaGap software (https://github.com/ViriatoII/MindaGap) using a tile size of 2144 a window size of 30 considering shifts calculated from transcripts with less than 400 copies in the window and occurring at least 10 times. The transcripts were then assigned to cells according to their overlap with the segmentation mask and analyzed as described below.

### Merscope

Tissues were sectioned in 10 µm slices and placed on one Merscope slide. Subsequently, the tissue was fixed with 4% PFA at 37°C for 15 min. After washing the with PBS, the sections were permeabilized with 70% ethanol at 4°C and until the hybridization was started. The panel (**Supplementary Dataset 1**) was hybridized for 48 hours, and all steps were performed according to the manufactures protocol including the membrane staining.

### Xenium

Tissues were sectioned in 10 µm slices and four samples were placed on one Xenium slide. Subsequently, the tissue was fixed with PFA according to the manufacturés protocol. Tissues were permeabilized with SDS, incubated in 70% ice cold methanol and washed with PBS. Hybridization of the human generic brain panel with 70 add-on genes (**Supplementary Dataset 1**) was performed at 50°C in a Bio-Rad C1000 touch cycler for 20 hours. Washing, ligation and amplification steps were carried out according to the manufacturer’s instructions. ROIs were selected according to the tissue area excluding non-tissue covered tiles. Each transcript was imaged in a bright state five times across 60 cycle-channels (15 cycles x 4 channels). After the run on the Xenium analyzer slides were removed and buffer exchanged with PBS-T for further storage at 4 °C.

### H&E staining

H&E staining of Visium slides was conducted by first removing the coverslip by incubation in 1x PBS Buffer followed by washing with H₂O. Next, slides were incubated in hematoxylin solution for 7 min and then washed with H₂O. Then, 300 µl bluing solution was added to the tissue, incubated for 2 min at room temperature and then washed in H₂O. Staining with an eosin solution (Sigma, 1:10 diluted in 0.45 M Tris acetic acid, pH=6) was performed for 1 min at room temperature followed by washing with H₂O. Then, slides were dehydrated by a series of washes at 70% (30 sec), 95% (30 sec) and two times 100% (1 min) ethanol and stored at room temperature. Virtual H&E staining followed the approach described previously [26]. Sections were stained with eosin solution for 1 min at room temperature, washed in H₂O and incubated for 15 min in 4x SSC (saline-sodium citrate) buffer. Sections were then stained with DAPI for 30 sec and mounted in Prolong Gold Antifade (Thermo Fisher Scientific). The H&E coloring of the DAPI and eosin staining was performed in R using the EBImage [46] together with custom scripts.

### Spinning disk confocal fluorescence microscopy

Imaging of RNAscope samples and reimaging of MC and Xenium slides by SDCM was conducted on an Andor Dragonfly 505 spinning disk confocal system equipped with a Nikon Ti2-E inverted microscope and a CFI P-Fluor 40X/1.30 oil objective or a Plan Apo 60x/1.40 oil objective. Multicolor images were acquired with the laser lines 405 nm (DAPI), 488 nm (Alexa 488, eosin), 561 nm (Atto 550), 637 nm (Atto 647) and 730nm (Alexa 750). Images were recorded at 16-bit depth and with 1024x1024 pixels dimensions (pixel size: 0.301 µm or 0.217 µm) using an iXon Ultra 888 EM-CCD camera. The region of interest was selected based on the DAPI signal and 50 z-slices were acquired with a step size of 0.4 µm (20 µm z-range) per field of view (FOV). Tiles were imaged with a 10% overlap to ensure accurate stitching. Subsequently, a flatfield-correction was conducted based on the DAPI channel and stitching and registration of the tiles was conducted with Fiji.

### Merscope amplification

Gene specific probes and amplification oligonucleotides were tested with the protein verification kit provided by Vizgen for the Merscope. A list of primary, secondary and amplification probes can be found in **Supplementary Table S6**. The tissue was fixed and permeabilized as described above, washed with 30% formamide in 2x SSC (wash buffer) and incubated with the primary probes at 1 µM concentration in hybridization buffer (0.05 % yeast rRNA, 1U/µl RNase inhibitor, 30% formamide, 2x SSC, 10% dextran). After 36 h of incubation at 37 °C in a humid environment the tissue was washed three times with wash buffer at 47 °C. The tissue was embedded according to the manufacturer’s instructions and incubated in clearing solution for 24h. Then, the tissue was washed with amplification buffer (10% formamide, 2x SSC) and the primary amplifier was hybridized at 5 nM in hybridization buffer (0.05 % yeast rRNA, 1U/µl RNase inhibitor, 10% formamide, 2x SSC, 10% dextran) for 30 min at 37°C. After three washes with amplification buffer the secondary amp probe was hybridized at 5 nM concentration in amp hybridization buffer (0.05 % yeast rRNA, 1U/µl RNase inhibitor, 10% formamide, 2x SSC, 10% dextran) for 30 min at 37 °C. After three washes the verification reagent was added for 15 min followed by a formamide and sample prep wash. The readout of the amplification probe was done with the protein verification kit (mouse, rabbit, goat) using only the mouse and rabbit channels.

### Sequential IF using the COMET platform

After completion of the Xenium run, the slides were washed twice with PBS and then placed in the comet system. The immuno-oncology SPYRE panel (**Supplementary Table S5**) was used to stain and image the tissue section of the sample MB299 using the standard SPYRE protocol on a Comet 1.0 instrument.

### Preprocessing of iST data for downstream analysis

For Xenium datasets (post XeniumRanger) we cropped selected areas since some tissue parts were folded/wrapped and disrupted. This is done to eliminate potential issues in further (downstream) analysis steps. A custom script is available at https://github.com/alikhuseynov/add-on_R/blob/develop/R/crop_seurat_v1.R and related discussion can be seen here https://github.com/satijalab/seurat/issues/8457.

### Cell segmentation

For cell segmentation the approach included in the Merscope (Cellpose 2 nuclei segmentation or cell segmentation with an additional cell boundary stain) and Xenium systems (nucleus segmentation with a custom neural network followed by a 15 µm Voronoi based cell boundary expansion) was used. For MC, cell segmentation was performed with Cellpose 2 as described above [8]. For an independent segmentation of the DAPI images of nuclei and cell membrane staining, if present, Cellpose 2 was used. Corresponding scripts to overlay images with the segmentation results were generated with the R script BrushUpSegmentationResults.R.

### Image processing and integration with ST data

Widefield images from the MC and Xenium platform were integrated with reimaged SDCM data with the following workflow in ImageJ. First, SDCM image stacks were subjected to a maximum intensity projection, followed by flat field and chromatic aberration correction using a custom script. Next, image tiles were stitched using the “Grid/Collection Stitching” plugin. DAPI images from SDCM were registered to MC or Xenium widefield images using “Register Virtual Stack Slices” with Affine feature extraction model and the Elastic bUnwarpJ splines registration model. In case of further staining, images were transformed via Transform Virtual Stack slices employing the transformation file of the DAPI registration.

### Combining data sets

Most of the analysis and visualization (including tidyverse, data.table, ggridges R packages) was done in R 4.2.2. Raw data were processed using technology-specific corporate pipelines (custom pipeline was used for MC). For each technology Seurat objects of the sample data and analysis results were created using the Seurat (v. 4.3.0) R package. For loading Vizgen/Merscope data and making a Seurat object, we optimized a loading function (see this PR https://github.com/satijalab/seurat/pull/7190), which was separately tested by Vizgen as well.

MC Seurat objects were created from the ROI file, segmentation mask, deduplicated transcripts and cell expression matrix generated with the resolve_processing pipeline (https://github.com/scOpenLab/resolve_processing, described above) with custom R scripts (https://github.com/scOpenLab/resolve-analysis). We merged technology-specific objects subset for same matching genes (96 in total) in a single object. When comparing to RNAscope only 10 matching genes were used. Cells with 0 counts were removed. To address issues with subset function on Seurat objects with spatial FOVs (see https://github.com/satijalab/seurat/issues/6409, https://github.com/satijalab/seurat/issues/ 7462) we wrote an optimized version https://github.com/scOpenLab/spatial_qc/blob/main/ scripts/subset_obj_seurat_v2.R, which was used in this study.

### Analysis of transcript counts per spatial bin or cell/nucleus

The distribution of transcript and feature/gene counts was analyzed for the shared set of 96 genes (**Supplementary Dataset 1**). It was either based on the number of transcripts in spatial bins with the size of a Visium spot of 55 µm diameter and 2,375 µm^2^ area, corresponding to a square side length of 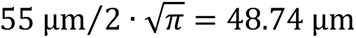, or on transcripts per nuclei/cell after segmentation. The spatial binning allows for an unbiased comparison at the selected bin size that is not confounded by effects of the cell/nucleus segmentation. At gene-level, we computed mean transcript counts across all cells and compared those value between the different technologies. Pairwise gene expression correlation analysis (Pearson correlation) within a cell was done for selected markers. The similarity to the RNAscope pattern was then computed as the coefficient of determination (R squared) of the correlation coefficients (**Supplementary Table S3**).

### Specificity analysis using background probes

To evaluate specificity of iST methods we used the probes included with the reagents for MC (25 false positive probes), Merscope (40 blank probes) and Xenium (128 unassigned codeword probes) (**Supplementary Dataset 1**) to which we here refer as background probes. Signals of 96 shared target genes and background were related based on their coordinates in a segmentation free manner. The number of target probes overlapping with background signal was determined by counting the spots of a given probe per tissue and ranking this sum of the probe signal. Averaged FDR values were calculated from the same data as

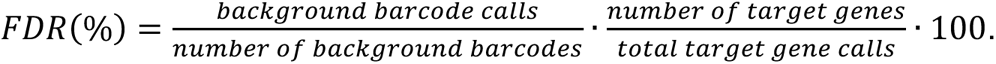

To evaluate the spatial distribution of target and background probes at the cell level, the spatial autocorrelation for each probe was computed as Moran’s *I* with the moranfast R package (C++ implementation). This function is similar to Moran.I from the ape R package but faster for large datasets. The input for computing Moran’s *I* with moranfast were the transcript counts per cell and the xy coordinates of cells centroids. The spatial neighborhood was defined using a distance-based (Euclidean) approach that computes distances *r* between pairs of cell centroids, this results in a distance matrix. The weighted inverse distance matrix was computed as 1 / distance matrix, the larger the resultant weight, the closer are the cell centroids. This approach was chosen over spatial contiguity-based approaches (queen, rook, hexagon, bishop spatial neighbors) since it does not require cell borders or polygons to touch each other. Bounds of Moran’s *I* go from -1 to +1 (similar to Pearson correlation coefficients). A value round 0 indicates spatially random pattern, < 0 towards -1 negative spatial autocorrelation (chessboard-like pattern), > 0 to towards 1 indicates positive spatial autocorrelation (clustered, also gradient-like patterns). This approach yields the spatial autocorrelation between transcripts at cellular resolution. Since our data set displayed no significant anticorrelation but only fluctuation around 0 (≥ -0.002) as their lower limit, we used min-max scaled Moran’s *I* from 0 to 1 in the plots shown.

As an alternative, molecule-level approach to assess spatial relations between the signal of a given probe, the distance to its nearest neighbor was calculated using the FNN R package with kd-tree search algorithm. The median of the resulting distribution was then used as the minimal distance value for further analysis (**Supplementary Dataset 2**).

### Integration, clustering and cell type annotation

We used Seurat SCTransform [47] and RunPCA to normalize data. Batch correction was performed using Harmony v1.0 [48] R package on samples (only two samples MB266 and 295) for each technology separately (MC, Xenium, Merscope), when integrating snRNAseq with those 3 spatial technologies, batch correction was also done on samples of the merged object (snRNAseq, MC, Xenium, Merscope). Clustering was performed for each technology on an integrated object using the Leiden algorithm [49] and visualized as UMAPs (all of these using Seurat). Cell type annotations were manually assigned according to the gene expression signatures reported previously [20].

## References

1. Sant P, Rippe K, Mallm JP: Approaches for single-cell RNA sequencing across tissues and cell types. Transcription 2023, 14:127–145.

2. Moffitt JR, Lundberg E, Heyn H: The emerging landscape of spatial profiling technologies. Nat Rev Genet 2022, 23:741–759.

3. Le P, Ahmed N, Yeo GW: Illuminating RNA biology through imaging. Nat Cell Biol 2022, 24:815–824.

4. Vandereyken K, Sifrim A, Thienpont B, Voet T: Methods and applications for single-cell and spatial multi-omics. Nat Rev Genet 2023, 24:494–515.

5. Moses L, Pachter L: Museum of spatial transcriptomics. Nat Methods 2022, 19:534–546.

6. Seferbekova Z, Lomakin A, Yates LR, Gerstung M: Spatial biology of cancer evolution. Nat Rev Genet 2023, 24:295–313.

7. Elhanani O, Ben-Uri R, Keren L: Spatial profiling technologies illuminate the tumor microenvironment. Cancer Cell 2023, 41:404–420.

8. Pachitariu M, Stringer C: Cellpose 2.0: how to train your own model. Nat Methods 2022, 19:1634–1641.

9. Petukhov V, Xu RJ, Soldatov RA, Cadinu P, Khodosevich K, Moffitt JR, Kharchenko PV: Cell segmentation in imaging-based spatial transcriptomics. Nat Biotechnol 2022, 40:345–354.

10. Greenwald NF, Miller G, Moen E, Kong A, Kagel A, Dougherty T, Fullaway CC, McIntosh BJ, Leow KX, Schwartz MS, et al: Whole-cell segmentation of tissue images with human-level performance using large-scale data annotation and deep learning. Nat Biotechnol 2022, 40:555–565.

11. Groiss S, Pabst D, Faber C, Meier A, Bogdoll A, Unger C, Nilges B, Strauss S, Föderl-Höbenreich E, Hardt M, et al: Highly resolved spatial transcriptomics for detection of rare events in cells. bioRxiv 2021:2021.2010.2011.463936.

12. Moffitt JR, Hao J, Bambah-Mukku D, Lu T, Dulac C, Zhuang X: High-performance multiplexed fluorescence in situ hybridization in culture and tissue with matrix imprinting and clearing. Proc Natl Acad Sci U S A 2016, 113:14456–14461.

13. Emanuel G, He J: Using MERSCOPE to Generate a Cell Atlas of the Mouse Brain that Includes Lowly Expressed Genes. Microscopy Today 2021, 29:16–19.

14. Lee H, Langseth CM, Salas SM, Metousis A, Alana ER, Garcia-Moreno F, Grillo M, Nilsson M: Open-source, high-throughput targeted in-situ transcriptomics for developmental biologists. bioRxiv 2023:2023.2010.2010.561689.

15. Janesick A, Shelansky R, Gottscho AD, Wagner F, Williams SR, Rouault M, Beliakoff G, Morrison CA, Oliveira MF, Sicherman JT, et al: High resolution mapping of the tumor microenvironment using integrated single-cell, spatial and in situ analysis. Nat Commun 2023, 14:8353.

16. Cook DP, Jensen KB, Wise K, Roach MJ, Dezem FS, Ryan NK, Zamojski M, Vlachos IS, Knott SRV, Butler LM, et al: A comparative analysis of imaging-based spatial transcriptomics platforms. bioRxiv 2023:2023.2012.2013.571385.

17. Wang H, Huang R, Nelson J, Gao C, Tran M, Yeaton A, Felt K, Pfaff KL, Bowman T, Rodig SJ, et al: Systematic benchmarking of imaging spatial transcriptomics platforms in FFPE tissues. bioRxiv 2023:2023.2012.2007.570603.

18. Salas SM, Czarnewski P, Kuemmerle LB, Helgadottir S, Matsson-Langseth C, Tismeyer S, Avenel C, Rehman H, Tiklova K, Andersson A, et al: Optimizing Xenium in situ data utility by quality assessment and best practice analysis workflows. bioRxiv 2023:2023.2002.2013.528102.

19. Hartman A, Satija R: Comparative analysis of multiplexed in situ gene expression profiling technologies. bioRxiv 2024:2024.2001.2011.575135.

20. Ghasemi DR, Okonechnikov K, Rademacher A, Tirier S, Maass KK, Schumacher H, Joshi P, Gold MP, Sundheimer J, Statz B, et al: Compartments in medulloblastoma with extensive nodularity are connected through differentiation along the granular precursor lineage. Nat Commun 2024, 15:269.

21. Northcott PA, Robinson GW, Kratz CP, Mabbott DJ, Pomeroy SL, Clifford SC, Rutkowski S, Ellison DW, Malkin D, Taylor MD, et al: Medulloblastoma. Nat Rev Dis Primers 2019, 5:11.

22. Korshunov A, Sahm F, Stichel D, Schrimpf D, Ryzhova M, Zheludkova O, Golanov A, Lichter P, Jones DTW, von Deimling A, et al: Molecular characterization of medulloblastomas with extensive nodularity (MBEN). Acta Neuropathol 2018, 136:303–313.

23. Wang H, Su N, Wang LC, Wu X, Bui S, Nielsen A, Vo HT, Luo Y, Ma XJ: Quantitative ultrasensitive bright-field RNA in situ hybridization with RNAscope. Methods Mol Biol 2014, 1211:201–212.

24. Wang F, Flanagan J, Su N, Wang LC, Bui S, Nielson A, Wu X, Vo HT, Ma XJ, Luo Y: RNAscope: a novel in situ RNA analysis platform for formalin-fixed, paraffin-embedded tissues. J Mol Diagn 2012, 14:22–29.

25. Moses L, Einarsson PH, Jackson K, Luebbert L, Booeshaghi AS, Antonsson S, Bray N, Melsted P, Pachter L: Voyager: exploratory single-cell genomics data analysis with geospatial statistics. bioRxiv 2023:2023.2007.2020.549945.

26. Lahiani A, Klaiman E, Grimm O: Enabling Histopathological Annotations on Immunofluorescent Images through Virtualization of Hematoxylin and Eosin. J Pathol Inform 2018, 9:1.

27. Simonson PD, Ren X, Fromm JR: Creating Virtual Hematoxylin and Eosin Images using Samples Imaged on a Commercial CODEX Platform. J Pathol Inform 2021, 12:52.

28. Xia C, Babcock HP, Moffitt JR, Zhuang X: Multiplexed detection of RNA using MERFISH and branched DNA amplification. Sci Rep 2019, 9:7721.

29. Okonechnikov K, Joshi P, Koerber V, Rademacher A, Bortolomeazzi M, Mallm J-P, da Silva PBG, Statz B, Sepp M, Sarropoulos I, et al: Medulloblastoma oncogene aberrations are not involved in tumor initiation, but essential for disease progression and therapy resistance. bioRxiv 2024:2024.2002.2009.579690.

30. Joshi P, Stelzer T, Okonechnikov K, Sarropoulos I, Sepp M, Pour-Jamnani MV, Rademacher A, Yamada-Saito T, Schneider C, Schmidt J, et al: Gene regulatory network landscape of Group 3/4 medulloblastoma. bioRxiv 2024:2024.2002.2009.579680.

31. Stickels RR, Murray E, Kumar P, Li J, Marshall JL, Di Bella DJ, Arlotta P, Macosko EZ, Chen F: Highly sensitive spatial transcriptomics at near-cellular resolution with Slide-seqV2. Nat Biotechnol 2021, 39:313–319.

32. Zhang Y, Petukhov V, Biederstedt E, Que R, Zhang K, Kharchenko PV: Gene panel selection for targeted spatial transcriptomics. Genome Biol 2024, 25:35.

33. Yafi MA, Hisham MHH, Grisanti F, Martin JF, Rahman A, Samee MAH: scGIST: gene panel design for spatial transcriptomics with prioritized gene sets. Genome Biol 2024, 25:57.

34. Schmidt U, Weigert M, Broaddus C, Myers G: Cell Detection with Star-convex Polygons. arXiv 2018:1806.03535.

35. Chen H, Li D, Bar-Joseph Z: SCS: cell segmentation for high-resolution spatial transcriptomics. Nat Methods 2023, 20:1237–1243.

36. Fu X, Lin Y, Lin DM, Mechtersheimer D, Wang C, Ameen F, Ghazanfar S, Patrick E, Kim J, Yang JYH: BIDCell: Biologically-informed self-supervised learning for segmentation of subcellular spatial transcriptomics data. Nat Commun 2024, 15:509.

37. Beštak K, Wuennemann F: nf-core/molkart: Spatial Circuit. Zenodo 2024:DOI: 10.5281/zenodo.10650749.

38. Blampey Q, Mulder K, Dutertre C-A, Gardet M, André F, Ginhoux F, Cournède P-H: Sopa: a technology-invariant pipeline for analyses of image-based spatial-omics. bioRxiv 2023:10.1101/2023.1112.1122.571863.

39. Cho CS, Xi J, Si Y, Park SR, Hsu JE, Kim M, Jun G, Kang HM, Lee JH: Microscopic examination of spatial transcriptome using Seq-Scope. Cell 2021, 184:3559–3572 e3522.

40. Chen A, Liao S, Cheng M, Ma K, Wu L, Lai Y, Qiu X, Yang J, Xu J, Hao S, et al: Spatiotemporal transcriptomic atlas of mouse organogenesis using DNA nanoball-patterned arrays. Cell 2022, 185:1777–1792 e1721.

41. Linares A, Brighi C, Espinola S, Bacchi F, Crevenna AH: Structured Illumination Microscopy Improves Spot Detection Performance in Spatial Transcriptomics. Cells 2023, 12.

42. Eng CL, Lawson M, Zhu Q, Dries R, Koulena N, Takei Y, Yun J, Cronin C, Karp C, Yuan GC, Cai L: Transcriptome-scale super-resolved imaging in tissues by RNA seqFISH. Nature 2019, 568:235–239.

43. Xia C, Fan J, Emanuel G, Hao J, Zhuang X: Spatial transcriptome profiling by MERFISH reveals subcellular RNA compartmentalization and cell cycle-dependent gene expression. Proc Natl Acad Sci U S A 2019, 116:19490–19499.

44. Bressan D, Battistoni G, Hannon GJ: The dawn of spatial omics. Science 2023, 381:eabq4964.

45. Zuiderveld K: Contrast limited adaptive histogram equalization. Graphics gems IV 1994:474–485.

46. Pau G, Fuchs F, Sklyar O, Boutros M, Huber W: EBImage - an R package for image processing with applications to cellular phenotypes. Bioinformatics 2010, 26:979–981.

47. Choudhary S, Satija R: Comparison and evaluation of statistical error models for scRNA-seq. Genome Biol 2022, 23:27.

48. Korsunsky I, Millard N, Fan J, Slowikowski K, Zhang F, Wei K, Baglaenko Y, Brenner M, Loh PR, Raychaudhuri S: Fast, sensitive and accurate integration of single-cell data with Harmony. Nat Methods 2019, 16:1289–1296.

49. Traag VA, Waltman L, van Eck NJ: From Louvain to Leiden: guaranteeing well-connected communities. Sci Rep 2019, 9:5233.

