## Supplementary figures and tables for "Comparison of spatial transcriptomics technologies using tumor cryosections"

### Content

#### Supplementary Figures

Figure S1. Workflow features of different ST methods.

Figure S2. Quantification of different imaging and segmentation methods.

Figure S3. Sensitivity analysis of iST methods.

Figure S4. Specificity analysis.

Figure S5. Clustering and cell type annotation.

#### Supplementary Tables

Table S1. Samples and readouts.

Table S2. Features of different *ST* methods.

Table S3. Similarity of marker gene correlation coefficient panel.

Table S4. Controls and their nomenclature used in *iST*.

Table S5. Antibody panel used with the COMET system.

Table S6. Primary, secondary and amplification probes for Merscope.

Table S7. Inventory of supplementary data sets associated with this manuscript.

Table S8. Inventory of datasets deposited at external repositories.

Table S9. Data analysis packages and software from external sources.

Table S10. Data analysis software from this study.

#### Supplementary Datasets

Additional datasets on samples and the analysis results derived from the different sequencing readouts have been uploaded as separate files in Microsoft Excel format with this manuscript (**Supplementary Table S7**) or have been deposited at external sources (**Supplementary Table S8**).

#### Supplementary References

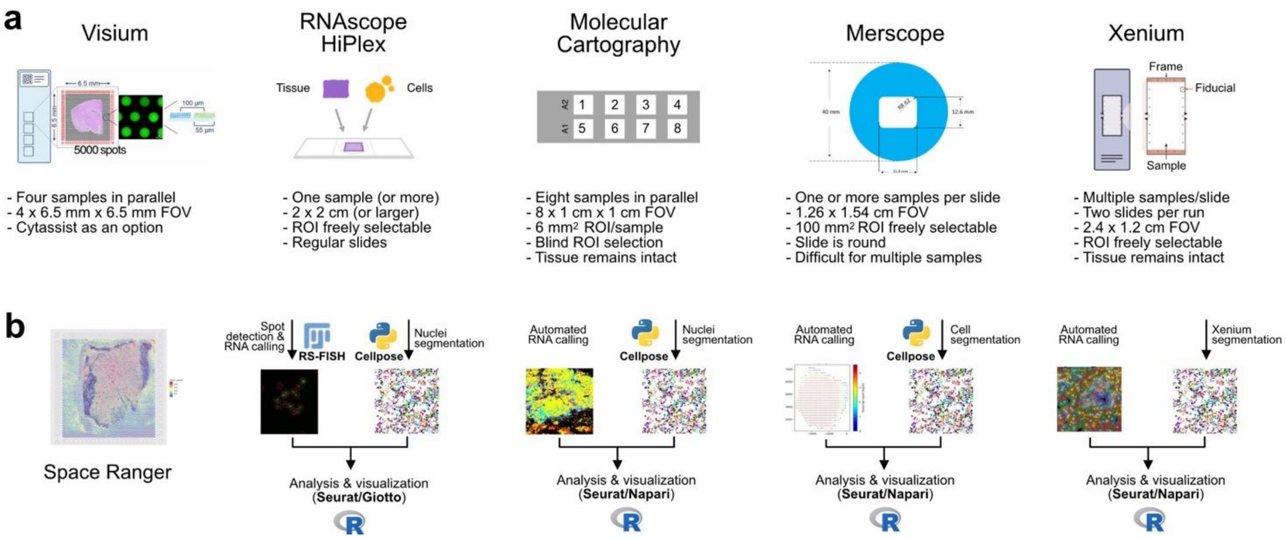


**Supplementary Figure 1. Workflow features of different ST methods.**

(**a**) Slide format and features. (**b**) Scheme of data analysis workflow used in the present study.

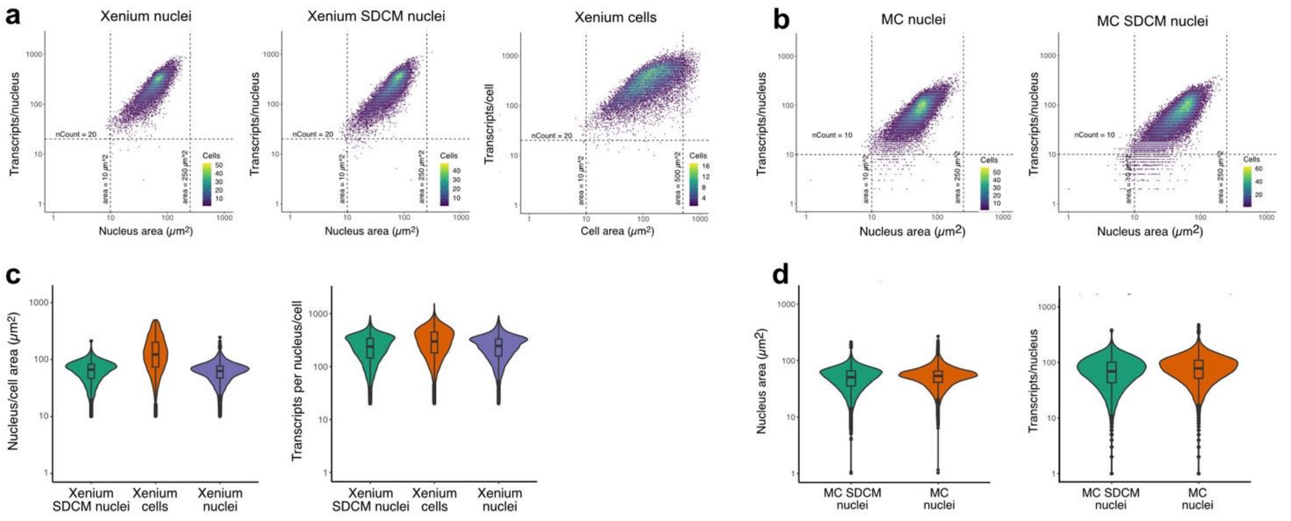


**Supplementary Figure 2. Quantification of different imaging and segmentation methods.**

(**a**) Quantification of the number of transcripts per nuclei/cell versus nuclei/cell area. The Xenium slide for an exemplary tumor tissue (MB266) was reimaged by SDCM (Xenium SDCM nuclei) and the segmentation results were compared with respect to the number of transcripts per nuclei/cell to the original widefield images acquired with Xenium analyzer by segmentation of nuclei with Cellpose (Xenium nuclei) of by using the Xenium segmentation workflow with expansion of the nuclei that aims to cover whole cells (Xenium cell). (**b**) Same as panel **a** but for Molecular Cartography (MC). (**c**) Violin plots showing the size distribution of segmented nuclei/cells (left) and transcripts per segmented nucleus/cell for MB266. SDCM refers to the spinning disk confocal images that are compared to widefield images acquired by the Xenium analyzer. (**d**) Same as panel **c** but for Molecular Cartography.

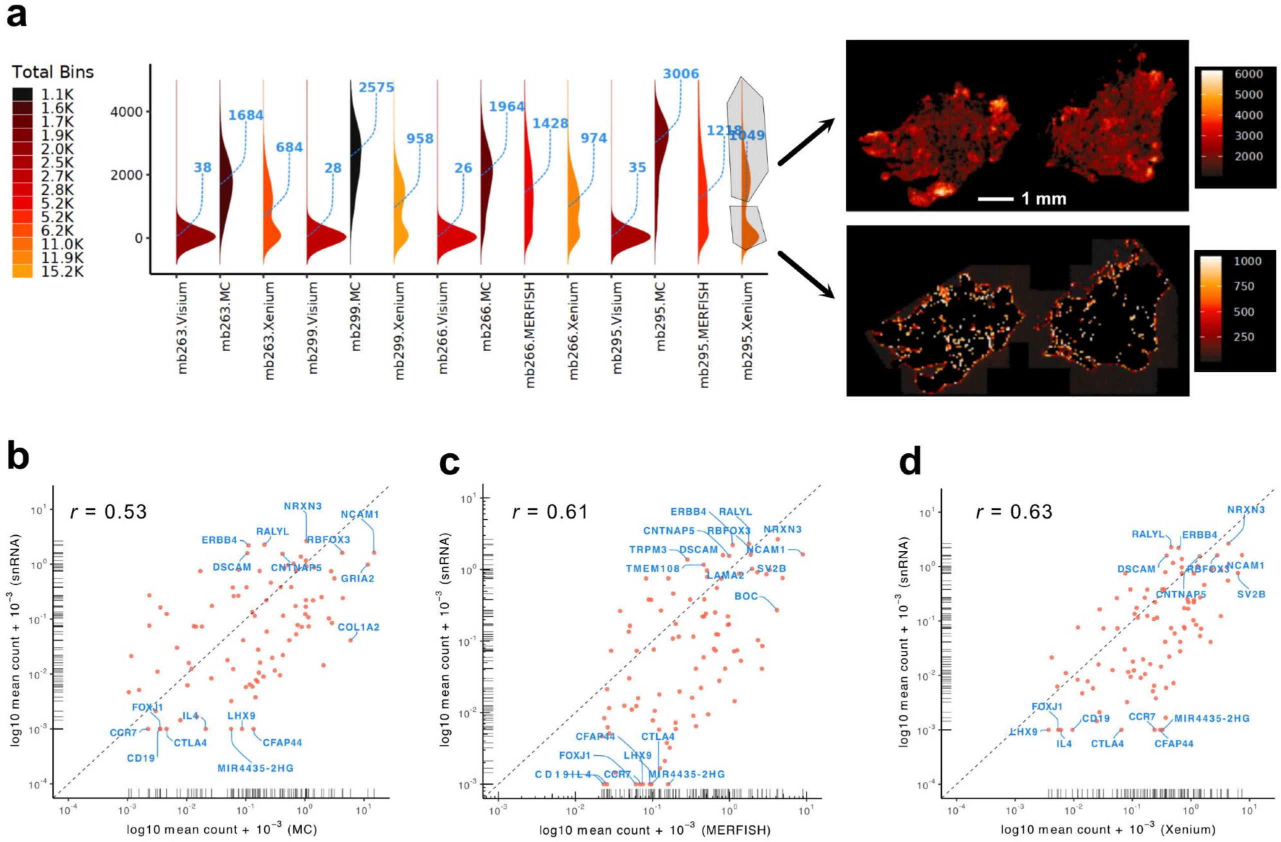


**Supplementary Figure 3. Sensitivity analysis of iST methods**

(**a**) Tissue location of cells in the high vs. low transcript distribution for the spatial binning analysis described in **Fig. 3a** for sample MB295 on Xenium. It can be seen that the cell population with the lower number of transcripts per bin is enriched in the outer regions of the tissue. (**b**) Correlation of snRNA-seq with MC for the shared panel of 96 genes snRNA-seq. The dashed line depicts the same number of transcripts detected for the two methods compared. The snRNA-seq analysis was conducted with the Chromium v2 chemistry for which a detection efficiency of 14-15% (v3 chemistry 30-32%) has been reported by the manufacturer (https://kb.10xgenomics.com/hc/en-us/articles/360001539051-What-fraction-of-mRNA-transcripts-are-captured-per-cell). (**c**) Same as panel b but for the correlation of snRNA-seq with Merscope. (**d**) Same as panel b but for the correlation of snRNA-seq with Xenium.

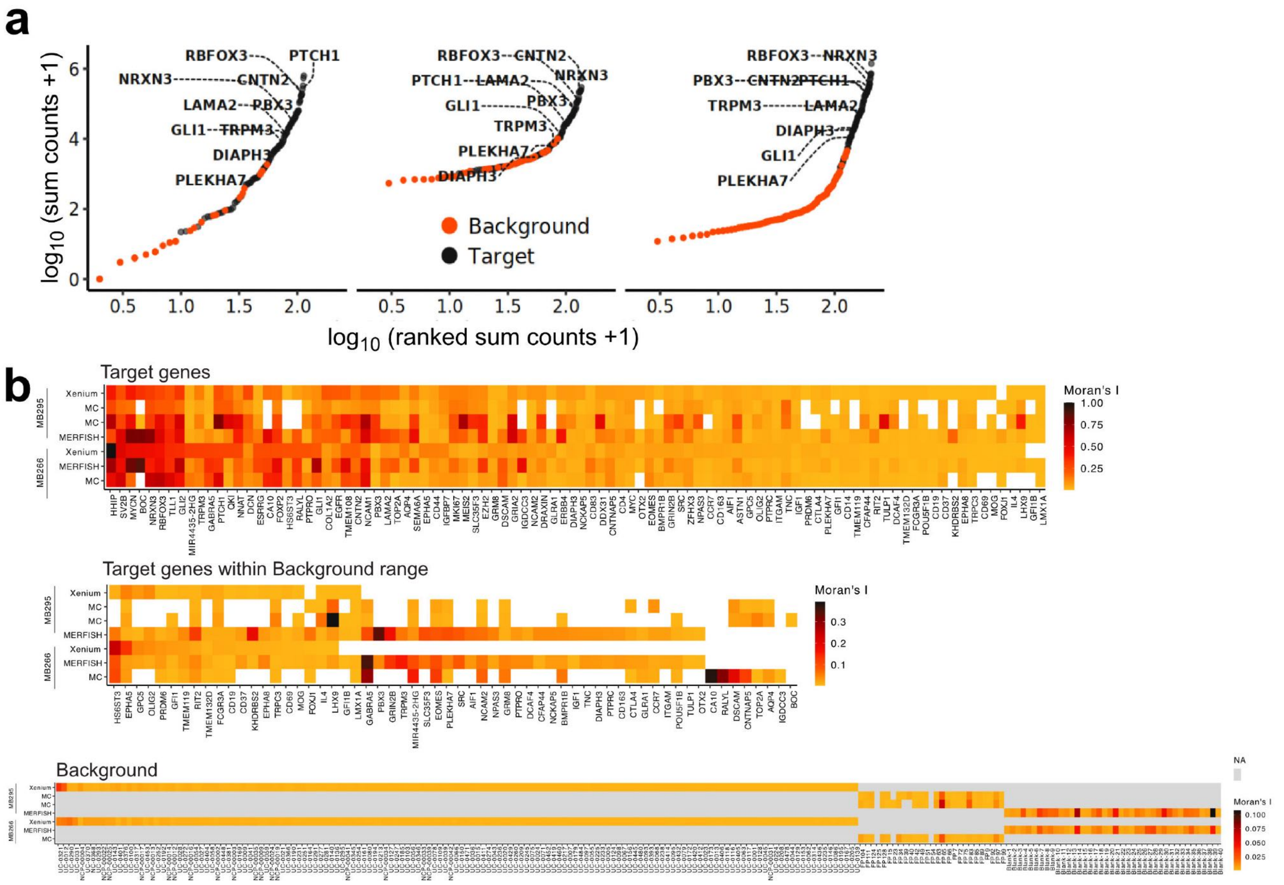


**Supplementary Figure 4. Specificity analysis.** The signal and spatial distribution of probes against the shared 96 RNAs ("target") was compared to the unspecific background signal. The corresponding control probes are called "false positive" (MC), "blank" (MERFISH) or "unassigned codeword and are referred to here as "background" (**Supplementary Table S4**). (**a**) Distribution of target and background signal plotted against the signal count for MBEN 295. (**b**) Spatial autocorrelation computed as Moran's *I* for target genes, target genes within background probes, and background probes (*I* is min-max scaled, range 0 to 1), p-value ≤ 0.05. A random and/or negative spatial autocorrelation is towards ~0.002 and 0. Abbreviations for Xenium background probes as described in **Table S4**: UC, unassigned codeword; NCP, negative control probe.

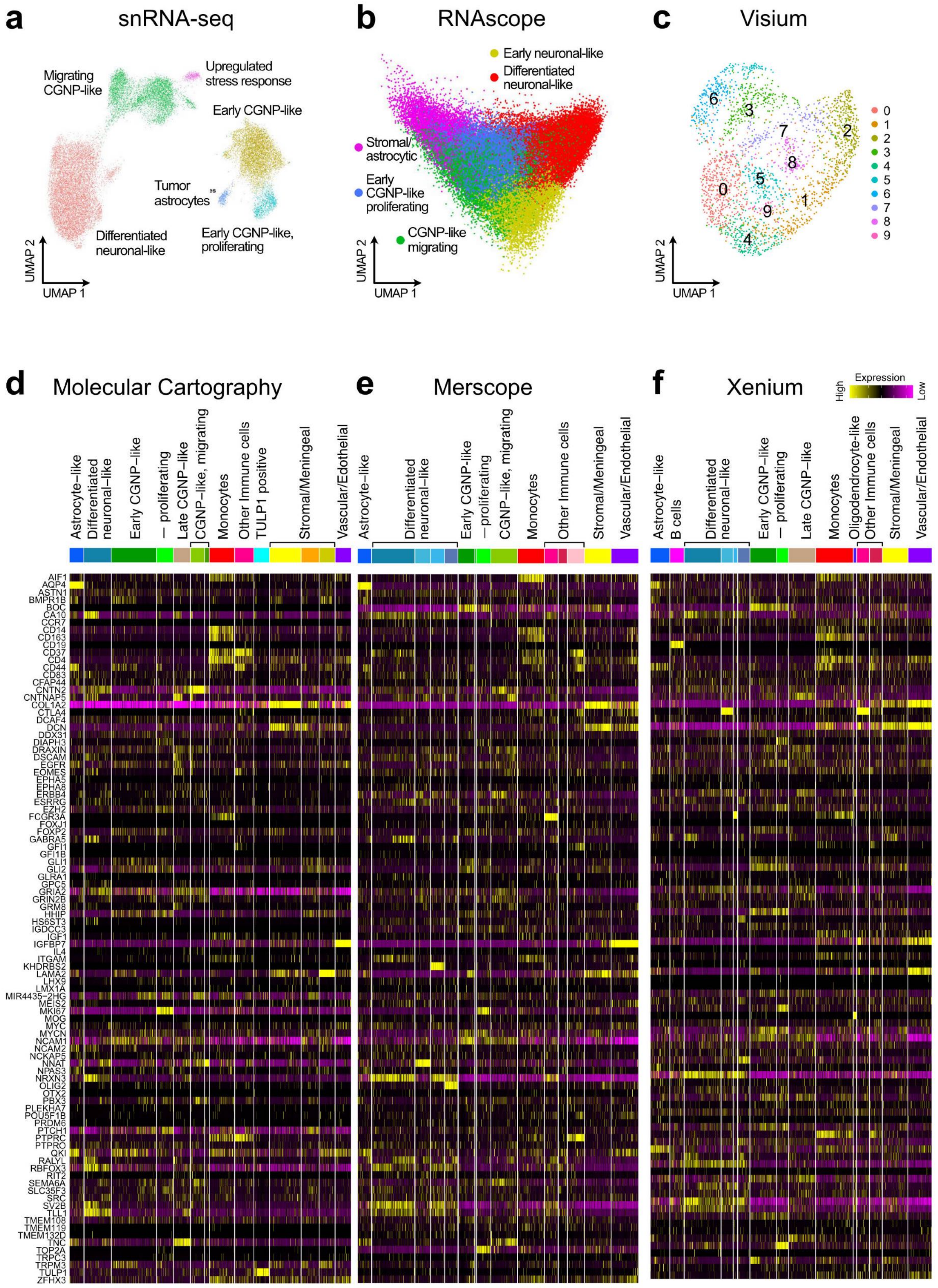


**Supplementary Figure 5. Clustering and cell type annotation.** Clustering was done with the panel of 96 shared genes without CD69, which was not a variable gene. (**a**) Clustering of scRNA-seq data for 27,782 cells taken from ref. [1]. (**b**) Clustering of RNAscope data for 110,508 cells taken from ref. [1]. (**c**) Visium. Due to insufficient spatial resolution, no cell types were assigned to the clusters obtained. (**d**) Heatmaps for clustering and cell type annotation of MC data. (**e**) Same as panel d but for Merscope. (**f**) Same as panel d but for Xenium.

**Table S1. Samples and readouts.**

| Sample ID | snRNA-seq | RNAscope | Visium | MC | Merscope | Xenium |
|---|---|---|---|---|---|---|
| MB263 | Yes | – | Yes | Yes | – | Yes |
| MB266 | Yes | Yes | Yes | Yes | Yes | Yes |
| MB295 | Yes | Yes | Yes | Yes | Yes | Yes |
| MB299 | Yes | Yes | Yes | Yes | – | Yes |

**Table S2. Features of different *ST* methods.**

| | Visium | RNAscope HiPlex | MC - Molecular Cartography | Merscope | Xenium |
|---|---|---|---|---|---|
| **Instrument requirements** | DNA sequencer | Fluorescence microscope (4-5 colors) | MC 1.0 | Merscope | Xenium Analyzer |
| **Resolution** | ~100 µm | subcellular | subcellular | subcellular | subcellular |
| **Gene panel number** | unbiased | 10 (shared) | 100 (96 shared with MERFISH & Xenium) | 138 (96 shared with MC) | 345 (96 shared with MC) |
| **Amplification** | PCR | ~100-1000 fluorophores per target | Direct labeling of RNA locus with >20 probes | Direct labeling of RNA locus with ~30-50 probes | Rolling circle amplification of padlock probes |
| **Transcript assignment** | 3'-sequencing & genome mapping | Fluorophore color | Combinatorial decoding in 8 imaging rounds | Combinatorial decoding in 16 imaging rounds | Combinatoria l decoding in 15 imaging rounds |
| **Controls** | – | Negative control targeting bacterial RNA | 25 FP (negative control codewords) | 40 blank (negative control codewords) | 128/41/20 unassingned /negative codewords/ negative probes |
| **H&E staining** | before run | after run | after run | not compatible | after run |

**Supplementary Table S3. Similarity of marker gene correlation coefficient panel.**

| | MC | MERFISH | Xenium | snRNA |
|---|---|---|---|---|
| **RNAScope** | 0.451 | 0.716 | 0.583 | 0.417 |
| **MC** | | 0.650 | 0.769 | 0.470 |
| **MERFISH** | | | 0.763 | 0.441 |
| **Xenium** | | | | 0.707 |

The correlation coefficients determined for the combinations of the 10 marker genes shown in **Fig. 4a** were compared between different technologies. The similarity was computed as R squared values of the correlation coefficients.

**Supplementary Table S4. Controls and their nomenclature used in *iST*.**

| Control | Origin | MC | Merscope | Xenium[d] | This study |
|---|---|---|---|---|---|
| **Secondary probe with code not in specific panel[a]** | Unspecific binding of fluorescently labeled secondary read-out probes | False positive | Blank | Unassigned codeword | Background probe |
| **Unspecific primary probe[b]** | Unspecific binding of random, non-targeting probe | Not used | Not used | Negative control probe | Negative control probe |
| **Unused decoding signal[c]** | Readout error that leads to an optical barcode that does not match that of a target gene | Not evaluated | Not evaluated | Negative control codeword | Not used |

The different technology platforms use different types of control probes for which the nomenclature and meaning is given in the table.

[a] The fluorescently labeled secondary probes with code sequence that are not present in the primary probe panel targeted against specific RNAs are called “false positive” for MC, “blank” (MERFISH) or “unassigned codeword” (Xenium). Here, they are referred to as background probes and represent false positive signal. The signal obtained with these probes can overlap with that of weakly expressed genes.

[b] The Xenium system uses additional controls with unspecific primary probes. Due to its amplification of the padlock probe signal, a single unspecifically bound probe can result in a target-like signal. With these controls a normalized negative control probe count $p_{neg}$ is calculated as $p_{neg} = p_{neg,total}/n_{neg}/n_{cells}$ where $p_{neg,total}$ is the total number of all negative control probe counts, $n_{neg}$ is the number of negative control probes) and $n_{cells}$ the number of cells. These estimated number of false positive transcripts per cell is then calculated as $p_{neg} \times n_{target}$ where $n_{target}$ is the number of target genes. Negative control probes are not needed for the MC and Merscope since they use 20-50 different probe per target. Here, the unspecific binding of a single primary probe would only lead to the binding one or two fluorophores. Thus, it would not be detectable unless the unspecific binding would occur at repetitive sequences.

[c] Target genes are identified from the optical barcodes acquired, i.e. the specific patterns of fluorescence signal in different colors generated by multiple rounds of secondary probe hybridization and imaging. Optical barcodes that do not correspond to that of a target represent errors in the readout of the fluorescence signals during image acquisition.

[d] In the Xenium system a quality score Q is assigned to each decoded transcript to assess the confidence in the decoded transcript identity. The Q value is derived from the likelihood of the maximum likelihood codeword (i.e., the codeword that best explains the observed data) compared to the likelihood of other sub-optimal codewords. Any target with a mean Q-score lower than 20 (Q20) is not included in further analysis.

**Supplementary Table S5. Antibody panel used with the COMET system.**

| Marker | Product code | Clone | Isotype | Host species | Recommended working conc. | | |
|---|---|---|---|---|---|---|---|
| | | | | | **in µg/ml** | **stock** | **Dilution** |
| αSMA | MR10100 | 1A4 | IgG/k | Mouse | 0.05 | 6.3 | 118 |
| CD3 | MR10010 | LUN3 | IgG1 | Rabbit | 0.14 | 241 | 1673 |
| CD4 | MR10020 | BL-155-1C11 | IgG | Rabbit | 10.00 | 1000 | 100 |
| CD8 | MR10030 | C8/144B | IgG1 | Mouse | 1.25 | 250 | 200 |
| CD11c | MR10070 | BLR138H | IgG | Rabbit | 0.67 | 100 | 150 |
| CD20 | MR10050 | L26 | IgG2a/k | Mouse | 0.11 | 35 | 312.5 |
| CD45 | MR10090 | PD7/26 + 2B11 | IgG1/k | Mouse | 7.71 | 993 | 128 |
| CD56 | MR10060 | LUN56 | IgG1 | Rabbit | 0.13 | 33 | 250 |
| CD68 | MR10080 | KP1 | IgG1 | Mouse | 0.20 | 50 | 250 |
| FOXP3 | MR10040 | 236A/E7 | IgG1 | Mouse | 19.10 | 955 | 50 |
| Ki-67 | MR10110 | BLR021E | IgG | Rabbit | 0.07 | 50 | 700 |
| PD-1 | MR10120 | EPR4877(2) | IgG | Rabbit | 2.50 | 2002 | 800 |
| PD-L1 | MR10130 | 73-10 | IgG | Rabbit | 0.14 | 115 | 800 |

**Supplementary Table S6. Primary, secondary and amplification probes for Merscope**.

| Target | Sequence |
|---|---|
| NES_primary_a | TGGGAGATTGAAGGTAGTGTTTCTTGAGGGGGTGGCCTCTGCTCTCCAGT GGTTAGAGTGAGTAGTAGTGGAGT |
| NES_primary_b | TGGGAGATTGAAGGTAGTGTTTTCTTCTTCTAGAGTCTTCAGTGGCTCCTG GTTAGAGTGAGTAGTAGTGGAGT |
| pre_amp_a_channel_2 | ACCCATTACTCCATTACCATATACCCATTACTCCATTACCATATACCCATTAC TCCATTACCATATACCCATTACTCCATTACCATATACCCATTACTCCATTACC ATATACTCCACTACTACTCACTCT |
| pre_amp_b_channel_2 | ACACTACCTTCAATCTCCCAAAACCCATTACTCCATTACCATATACCCATTA CTCCATTACCATATACCCATTACTCCATTACCATATACCCATTACTCCATTAC CATATACCCATTACTCCATTACCAT |
| sec_amp_channel_2 | TTGTAAATGGAGGTGGTATATTGTAAATGGAGGTGGTATATTGTAAATGGAG GTGGTATATTGTAAATGGAGGTGGTATAATGGTAATGGAGTAATGGGT |

**Supplementary Table S7. Inventory of supplementary datasets associated with this study.**

| File Name | Figure/Table | Description |
|---|---|---|
| Supplementary Dataset 1: SuppDataset01.xlsx | Fig. 1; Table 1; | Probe set used for the different iST methods. |
| Supplementary Dataset 2: SuppDataset02.xlsx | Fig. 4 | Expression levels, spatial autocorrelation, and median nearest neighbor distance. |
| Supplementary Dataset 3: SuppDataset03.xlsx | Fig. 5, Fig. S5 | Expression signatures and cell type annotation of MC, Merscope and Xenium data. |

**Supplementary Table S8. Inventory of datasets deposited at external repositories.**

| Data | Repository | Link | Description |
|---|---|---|---|
| snRNA-seq count tables | GEO, GSE239854 | https://www.ncbi.nlm.nih.gov/geo/query/acc.cgi?acc=GSE239854 | Data for samples MB266, MB295, MB299 from ref. [1]. |
| MC transcript count tables | GEO, GSE247736 | https://www.ncbi.nlm.nih.gov/geo/query/acc.cgi?acc=GSE247736 | Data for samples MB263, MB266, MB295, MB299 from ref. [1] |
| RNAscope | BioImage Archive, S-BIAD826 | https://www.ebi.ac.uk/biostudies/bioimages/studies/S-BIAD826 | Raw and processed data from ref. [1] |
| MC (Molecular Cartography) | BioImage Archive, S-BIAD825 | https://www.ebi.ac.uk/biostudies/bioimages/studies/S-BIAD825?query=S-BIAD825 | Raw and processed data for samples MB263, MB266, MB295, MB299 from ref. [1] |
| Visium | Zenodo | https://doi.org/10.5281/zenodo.10863259 | Raw and filtered feature bc matrix and spatial data for samples MB263, MB266, MB295, MB299, this study. |
| MC (additional data), Merscope, Xenium | BioImage Archive, S-BIAD1093 | https://www.ebi.ac.uk/biostudies/bioimages/studies/S-BIAD1093?query=S-BIAD1093 | Raw and processed data for samples MB263, MB266, MB295, MB299, this study. |
| Seurat objects for cell-based analysis | Zenodo | https://doi.org/10.5281/zenodo.10863259 | Seurat objects for cell-based analysis of shared set of 96 genes for snRNA-seq, Merscope, MC and Xenium for and for shared set of 10 genes when including RNAscope, this study. |
| Seurat objects for segmentation-free analysis | Zenodo | https://doi.org/10.5281/zenodo.10863259 | Seurat objects for segmentation-free analysis of shared set of 96 genes for snRNA-seq, Merscope, MC and Xenium for and for shared set of 10 genes when including RNAscope, this study. |

**Supplementary Table S9. Data analysis packages and software from external sources.**

| Software | Ref. | Link | Version |
|---|---|---|---|
| Bioconductor | [2] | www.bioconductor.org | 1.30.18 |
| Seurat | [3] | satijalab.org/seurat/ | 4.3.0.9002 |
| Cellpose 2 | [4] | https://www.cellpose.org/ | v2.1.1 |
| Cellpose | [5] | https://pypi.org/project/cellpose/ | 2.2 |
| Voyager | [6] | https://pachterlab.github.io/voyager/ | |
| Harmony | [7] | https://cran.r-project.org/package=harmony | 1.0 |
| SCTransform | [2] | https://cran.r-project.org/package=sctransform | |
| roifile | [8] | https://pypi.org/project/roifile/ | 2023.2.12 |
| Pillow | [9] | https://python-pillow.org/ | 9.4.0 |
| tifffile | [10] | https://pypi.org/project/tifffile/ | v2023.3.15 |
| scikit-image | [11] | https://scikit-image.org/ | 0.2.0 |
| opencv-python-headless | [12] | https://pypi.org/project/opencv-python/ | 4.7.0.72 |
| QuPath | [13] | https://qupath.github.io/ | 0.5.0 |
| sf (simple feature access) | [14] | https://r-spatial.github.io/sf/ | 1.0-15 |
| RImageJROI | | https://cran.r-project.org/package=RImageJROI | 0.1.2 |
| geojsonsf | | https://cran.r-project.org/web/packages/geojsonsf/index.html | 2.0.3 |
| moranfast | | https://github.com/mcooper/moranfast | … |

**Supplementary Table S10. Custom data analysis software from this study.**

| Software | Link | Description |
|---|---|---|
| spatial_qc | https://github.com/scOpenLab/spatial_qc | R scripts for QC of ST data. |
| spatial_analysis | https://github.com/scOpenLab/spatial_analysis | R-scripts and Jupyter notebooks for analysis of ST data. |
| resolve_processing | https://github.com/scOpenLab/resolve_processing | Tools and scripts for analyzing MC data with a Nextflow pipeline for segmentation and computing transcripts counts per cell |
| seurat_resolve_importer | https://github.com/scOpenLab/seurat_resolve_importer | Tools and scripts for analyzing MC data with Seurat. |
| geojson_seurat_cropper | https://github.com/scOpenLab/geojson_seurat_cropper | Crops a Seurat object using a sf polygon to retain only cells and molecules inside the polygon. |
| xenium_processing | https://github.com/scOpenLab/xenium_processing | Scripts for processing of Xenium data. |
| R scripts for reimaging and image analysis | https://github.com/RippeLab/MBEN | Scripts used for image processing, image registration and downstream data analysis of RNAscope, MC and Xenium data after reimaging. |

### Supplementary References


1. Ghasemi DR, Okonechnikov K, Rademacher A, Tirier S, Maass KK, Schumacher H, Joshi P, Gold MP, Sundheimer J, Statz B, et al: **Compartments in medulloblastoma with extensive nodularity are connected through differentiation along the granular precursor lineage.** *Nat Commun* 2024, **15:**269.

2. Gentleman RC, Carey VJ, Bates DM, Bolstad B, Dettling M, Dudoit S, Ellis B, Gautier L, Ge Y, Gentry J, et al: **Bioconductor: open software development for computational biology and bioinformatics.** *Genome Biol* 2004, **5:**R80.

3. Stuart T, Butler A, Hoffman P, Hafemeister C, Papalexi E, Mauck WM, 3rd, Hao Y, Stoeckius M, Smibert P, Satija R: **Comprehensive Integration of Single-Cell Data.** *Cell* 2019, **177:**1888-1902 e1821.

4. Pachitariu M, Stringer C: **Cellpose 2.0: how to train your own model.** *Nat Methods* 2022, **19:**1634-1641.

5. Moses L, Einarsson PH, Jackson K, Luebbert L, Booeshaghi AS, Antonsson S, Bray N, Melsted P, Pachter L: **Voyager: exploratory single-cell genomics data analysis with geospatial statistics.** *bioRxiv* 2023**:**2023.2007.2020.549945.

6. Korsunsky I, Millard N, Fan J, Slowikowski K, Zhang F, Wei K, Baglaenko Y, Brenner M, Loh PR, Raychaudhuri S: **Fast, sensitive and accurate integration of single-cell data with Harmony.** *Nat Methods* 2019, **16:**1289-1296.

7. Choudhary S, Satija R: **Comparison and evaluation of statistical error models for scRNA-seq.** *Genome Biol* 2022, **23:**27.

8. Gohlke C: **cgohlke/roifile: v2023.2.12.** *Zenodo* 2023**:**doi: 10.5281/zenodo.7633998.

9. Murray A, Kemenade Hv, wiredfool, Clark JA, Alexander Karpinsky, Baranovič O, Gohlke C, Dufresne J, DWesl, Schmidt D, et al: **python-pillow/Pillow: 9.4.0.** *Zenodo* 2023**:**doi: 10.5281/zenodo.7498081.

10. Gohlke C: **cgohlke/tifffile: v2023.3.15.** *Zenodo* 2023**:**doi: 10.5281/zenodo.7738996.

11. van der Walt S, Schonberger JL, Nunez-Iglesias J, Boulogne F, Warner JD, Yager N, Gouillart E, Yu T, scikit-image c: **scikit-image: image processing in Python.** *PeerJ* 2014, **2:**e453.

12. Bradski G: **The OpenCV Library. Dr. Dobb's Journal of Software Tools.** *Dr Dobb's J Softw Tools* 2000, **120:**122-125.

13. Bankhead P, Loughrey MB, Fernandez JA, Dombrowski Y, McArt DG, Dunne PD, McQuaid S, Gray RT, Murray LJ, Coleman HG, et al: **QuPath: Open source software for digital pathology image analysis.** *Sci Rep* 2017, **7:**16878.

14. Pebesma E, Bivand R: *Spatial Data Science: With Applications in R.* 1 edn: Chapman and Hall/CRC; 2023.
